# Human iPSC gene signatures and X chromosome dosage impact response to WNT inhibition and cardiac differentiation fate

**DOI:** 10.1101/644633

**Authors:** Agnieszka D’Antonio-Chronowska, Margaret K. R. Donovan, Paola Benaglio, William W. Greenwald, Michelle C. Ward, Hiroko Matsui, Kyohei Fujita, Sherin Hashem, Francesca Soncin, Mana Parast, Eric Adler, Erin N. Smith, Matteo D’Antonio, Kelly A. Frazer

## Abstract

Non-genetic variability in human induced pluripotent stem cell (iPSC) lines impacts their differentiation outcome, limiting their utility for genetic studies and clinical applications. Despite the importance of understanding how non-genetic molecular variability influences iPSC differentiation outcome, large-scale studies capable of addressing this question have not yet been conducted. Here, we performed 258 directed differentiations of 191 iPSC lines using established protocols to generate iPSC-derived cardiovascular progenitor cells (iPSC-CVPCs). We observed cellular heterogeneity across the iPSC-CVPC samples due to varying fractions of two cell types: cardiomyocytes (CMs) and epicardium-derived cells (EPDCs). Analyzing the transcriptomes of CM-fated and EPDC-fated iPSCs discovered that 91 signature genes and X chromosome dosage differences influence WNT inhibition response during differentiation and are associated with cardiac fate. Analysis of an independent set of 39 iPSCs differentiated to the cardiac lineage confirmed shared sex and transcriptional differences that impact cardiac fate outcome. The scale and systematic approach of our study enabled novel insights into how iPSC transcriptional and X chromosome gene dosage differences influence WNT signaling during differentiation and hence cardiac cell fate.

## Introduction

Variability in human induced pluripotent stem cell (iPSC) lines compromises their utility for regenerative medicine and as a model system for genetic studies. This variability impacts iPSC differentiation outcome and despite using standardized differentiation protocols, large-scale applications result in the generation of samples with cellular heterogeneity (i.e. multiple cell types are present within a given sample and the proportions of cell types vary across samples) (Guzzo et al., 2013; Schwartzentruber et al., 2018; Zhao et al., 2017). Previous large-scale quantitative trait loci (QTL) studies in iPSCs (Carcamo-Orive et al., 2017; DeBoever et al., 2017; Kilpinen et al., 2017) have shown that genetic variation accounts for the majority of expression differences between iPSC lines, but non-genetic factors also contribute to these differences. Understanding how non-genetic transcriptional differences between iPSC lines impact their differentiation outcome is necessary to improve the ability to generate cell types of interest.

Well-established small molecule protocols for generating iPSC-derived cardiovascular progenitor cells (iPSC-CVPCs) (Lian et al., 2013) produce fetal-like cardiomyocytes, which can undergo further specification as cells mature in culture into various cardiac subtypes (atrial, ventricular, or nodal) (Burridge et al., 2014). Based on variable cardiac troponin T (cTnT) staining, the derived samples are known to display cellular heterogeneity (Dubois et al., 2011; Kattman et al., 2011; Witty et al., 2014), but the origin of the cTnT-negative non-myocyte cells, and whether the same or different non-myocyte cell types are consistently derived alongside cTnT-positive myocytes across samples has not previously been investigated. The differentiation protocol is dependent on manipulation of WNT signaling, initially through activation of the pathway by GSK3 inhibition, followed by inhibition of the pathway by Porcupine (PORCN) inhibition (Mo et al., 2013; Wang et al., 2013). An in-depth analysis of the outcomes of independent differentiations of hundreds of iPSC lines with different genetic backgrounds could provide insights into the origins of the non-myocyte cells, as well as the extent to which non-genetic transcriptional differences between iPSC lines contribute to the iPSC-CVPC cellular heterogeneity.

Here, we used a highly standardized and systematic approach to conduct 258 directed differentiations of 191 iPSC lines (Panopoulos et al., 2017) into iPSC-CVPCs. We characterized the cellular heterogeneity of the iPSC-CVPC samples and showed that only two distinct cell types were present, cardiomyocytes (CMs) and epicardium-derived cells (EPDCs), which varied in proportion across samples. As differentiation protocols to derive iPSC-CMs and iPSC-EPDCs primarily differ by a step involving WNT inhibition to derive the former, but not the later (Bao et al., 2016; Hartman et al., 2016; Iyer et al., 2015; Witty et al., 2014), we hypothesized that the observed cellular heterogeneity could result from suboptimal WNT inhibition in subsets of cells across iPSC lines. To test this hypothesis, we analyzed transcriptional differences between iPSC lines that differentiated into CMs and those that differentiated into EPDCs (e.g. iPSCs with a CM-fate or EPDC-fate) and discovered 91 signature genes associated with cardiac fate differentiation outcome. These signature genes are involved in differentiation, including the Wnt/β-catenin pathway, muscle differentiation or cardiac-related functions, and the transition of epicardial cells to EPDCs by epithelial-mesenchymal transition (EMT). While the proportion of variance explained by each of the signature genes varied over three orders of magnitude, altogether they captured approximately half of the total variance underlying iPSC fate determination. Additionally, we show variability in X chromosome gene dosage (X_active_X_active_ vs X_active_X_inactiv_e vs XY) across iPSCs plays a role in cardiac fate determination. The association with X chromosome gene dosage could in part be due to higher expression in CM-fated iPSCs of chrXp11 genes, which encodes *PORCN.* Transcriptomic analysis of an independent set of 39 iPSCs differentiated to the cardiac lineage using a similar small molecule protocol (Banovich et al., 2018) confirmed our findings.

## Results

### iPSC-CVPCs show cellular heterogeneity across samples

To gain insights into molecular mechanisms that could influence variability in human iPSC differentiation outcome, we employed a highly systematic approach (Figures S1, S2) to differentiate 191 pluripotent lines from 181 iPSCORE individuals (Panopoulos et al., 2017) (Figure 1A, B, Table S1) into iPSC-derived cardiovascular progenitor cells (iPSC-CVPCs). We used a small molecule cardiac differentiation protocol used to derive cardiomyocytes (Lian et al., 2015) followed on D15 by lactate selection to obtain pure cardiac cells (Tohyama et al., 2013). In total, we conducted 258 differentiations, of which 193 (80.6%, from 154 lines derived from 144 subjects) were completed, i.e. reached Day 25 of differentiation, while 65 (from 37 lines derived from 37 subjects) were terminated prior to Day 25, because they did not form a syncytial beating monolayer (Table S2, Table S3). The completed iPSC-CVPCs at D25 on average had a high fraction of cells that stained positive for cardiac troponin T (%cTnT, median = 89.2%; Figure 1C) and were positive by immunofluorescence (IF) for cardiac markers (Figure 1D-G, Figure S3, Table S4); however, 15 lines had %cTnT < 40%, indicating that despite lactate selection, there was substantial cellular heterogeneity within and across samples.

**Figure 1:**
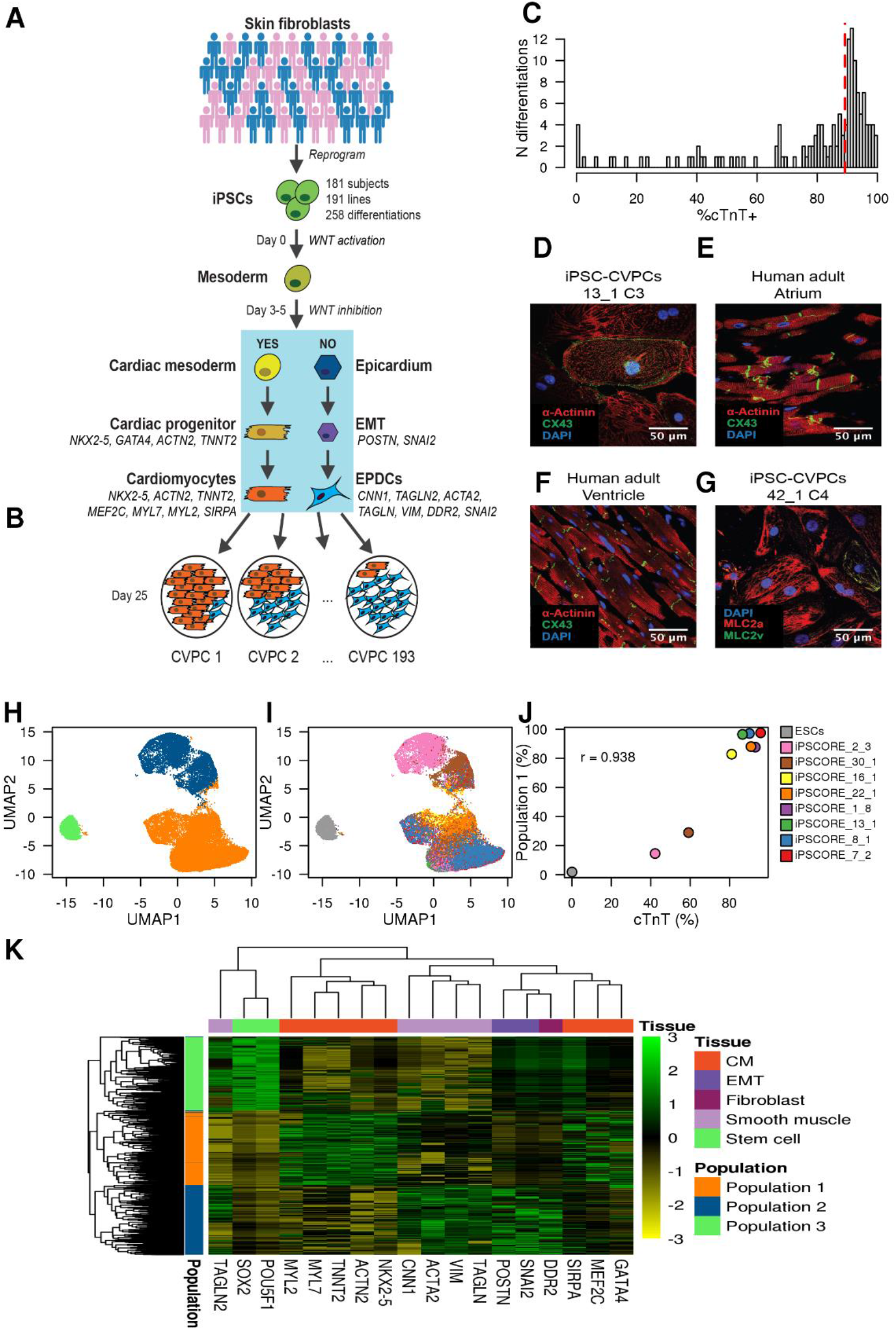
Characterization of cellular heterogeneity in iPSC-CVPC samples. (A, B) Overview of the study design. (A) Skin fibroblasts from 181 subjects were reprogrammed to iPSCs and differentiated to iPSC-CVPCs (191 lines, 258 differentiations). After WNT pathway activation at day 0 (D0) and its inactivation by IWP-2 at D3-5, cells differentiate to cardiomyocytes (CMs) if the WNT signaling is successfully inhibited. If WNT signaling is not sufficiently inhibited, cells differentiate to EPDCs. (B) 193 of the 258 differentiations were completed (D25), and we observed that different CVPC samples had different proportions of CMs and EPDCs. (C) Distribution of %cTnT. Dashed red line represents the median value. (D-F) Immunofluorescence staining of (D) iPSC-CVPCs, (E) human atrium, and (F) ventricle with IF markers DAPI (blue), ACTN1 (red), and CX43 (green). (G) Immunofluorescence staining iPSC-CVPCs with IF markers DAPI (blue), MLCa+ (red) and MLCv+ (green). (H) scRNA-seq UMAP plot showing the presence of three populations: CMs (orange), EPDCs (blue) and ESCs (green). (I) scRNA-seq UMAP plot showing the distribution of the nine analyzed samples (8 iPSC-CVPC lines and one ESC line) across the different clusters. (J) Scatterplot showing the correlation between the %cTnT and the fraction of cells in Population 1 (CMs) for each of the nine samples. (K) Heatmap showing across all 34,905 single cells the expression markers for: 1) stem cells *(POU5F1; SOX2);* 2) CMs *(GATA4, MEF2C, SIRPA, TNNT2, MYL7, ACTN2, NKX2-5, MYL2);* 3) EMT *(SNAI2, POSTN);* 4) fibroblasts (DDR2); and 5) smooth muscle *(TAGLN2, CNN1, ACTA2, VIM, TAGLN).*

### Subset of cells show differential response to WNT inhibition during differentiation

To examine the cellular heterogeneity in the iPSC-CVPCs, we performed single-cell RNA-seq (scRNA-seq) on eight samples with varying %cTnT values (42.2 to 95.8%, Table S5, Table S6) and combined these data with scRNA-seq from the H9 ESC line (total of 34,905 cells). We detected three distinct cell populations: 1) Population 1, 21,056 cells (60.3%); 2) Population 2, 11,044 cells (31.6%); and 3) Population 3, 2,805 cells (8.1%) (Figure 1H, Figure S4), (Table S7). While Populations 1 and 2 were comprised of the eight iPSC-derived samples, Population 3 was almost exclusively included ESC cells (97.7% of the 2,870 ESC cells), (Figure 1I, Figure S5). The relative proportion of cells that each of the iPSC-CVPC samples contributed to Population 1 versus Population 2 was strongly correlated with its %cTnT value (r = 0.938, p = 1.89 × 10^−4^, t-test; Figure 1J), suggesting that Population 1 was cardiomyocytes.

As cardiomyocytes (CMs) and epicardium lineage cells could both survive lactate purification (Friedman et al., 2018; Iyer et al., 2015; Ng et al., 1997; Tohyama et al., 2013), we investigated if the non-myocyte cells composing Population 2 were iPSC-epicardium-derived cells (iPSC-EPDCs). We examined the expression levels of 16 marker genes (Figure 1A, B) specific for either CMs or EPDCs (including smooth muscle, fibroblasts and genes involved in epithelial-to-mesenchymal transition, Figure 1K) and two marker genes for stem cells. Consistent with having a high number of cTnT-positive cells, Population 1 expressed high levels of CM-specific genes, while Population 2 expressed high levels of EPDC-specific genes, and Population 3 expressed high levels of the stem cell markers *POU5F1* and *SOX2* (Figure S6). Of note, *TNNT2* was expressed in some of the cells in Population 2 (Figure S6), which is consistent with the strong, but not absolute correlation between %cTnT value and fraction of Population 1 (Figure 1J), and previous studies showing that some EPDCs express *TNNT2* (Witty et al., 2014). These results show that the small molecule differentiation protocol followed by lactate purification (Figure S1) resulted in the absence of undifferentiated cells at D25 and in the derivation of two distinct cell populations, one of which expresses high levels of CM markers, including *TNNT2, NKX2-5* and *MEF2C* (Population 1), and the other which expresses EPDC markers, including *SNAI2, DDR2, VIM* and *ACTA2* (Population 2). Of note, the protocols for generating iPSC-derived cardiomyocytes (iPSC-CMs) and iPSC-EPDCs both involve activating the WNT signaling pathway (Bao et al., 2016; Iyer et al., 2015; Witty et al., 2014) and have a shared intermediate mesoderm progenitor, but subsequent WNT inhibition directs differentiating cells to iPSC-CMs and endogenous levels of WNT signaling direct differentiating cells to iPSC-EPDCs (Witty et al., 2014) (Figure 1A). Therefore, our results suggest that iPSC-CVPC cellular heterogeneity results from suboptimal WNT inhibition in a subset of cells during differentiation, which then give rise to EPDCs.

### iPSC-CVPCs are composed of immature CMs and EPDCs

To estimate the relative abundances of CM and EPDC cells across our collection of iPSC-CVPC samples, we selected the top 50 significantly overexpressed genes in each of the three scRNA-seq populations (150 genes in total, p < 10^−13^, edgeR, Table S8), obtained their expression levels in bulk RNA-seq from 180 iPSC-CVPCs (Table S5), and inputted these values into CIBERSORT (Newman et al., 2015). We observed that the proportions of each cell type varied across the samples, although the iPSC-CVPCs tended to have a greater fraction of CMs (84.8 ± 31.8%, Figure 2A) than EPDCs (14.7 ± 32.0%), and essentially no stem cells (0 ± 0. 8%). The estimated fraction of CMs and EPDCs in the iPSC-CVPCs was highly correlated with %cTnT values (r = 0.927, p ≈ 0; t-test Figure 2B), similar to that observed in the analysis of the scRNA-seq data (Figure 1J). Finally, we showed that the iPSC-CVPCs with high estimated CM or EPDC cellular fractions respectively showed higher expression of CM markers *(MEF2C, NKX2-5* and *ACTN2)* and EPDC markers *(ACTA2, TAGLN, DDR2* and *SNAI2)* (Figure 2C). These results indicate that cellular heterogeneity across iPSC-CVPC samples largely reflects different proportions of CMs and EPDCs.

**Figure 2:**
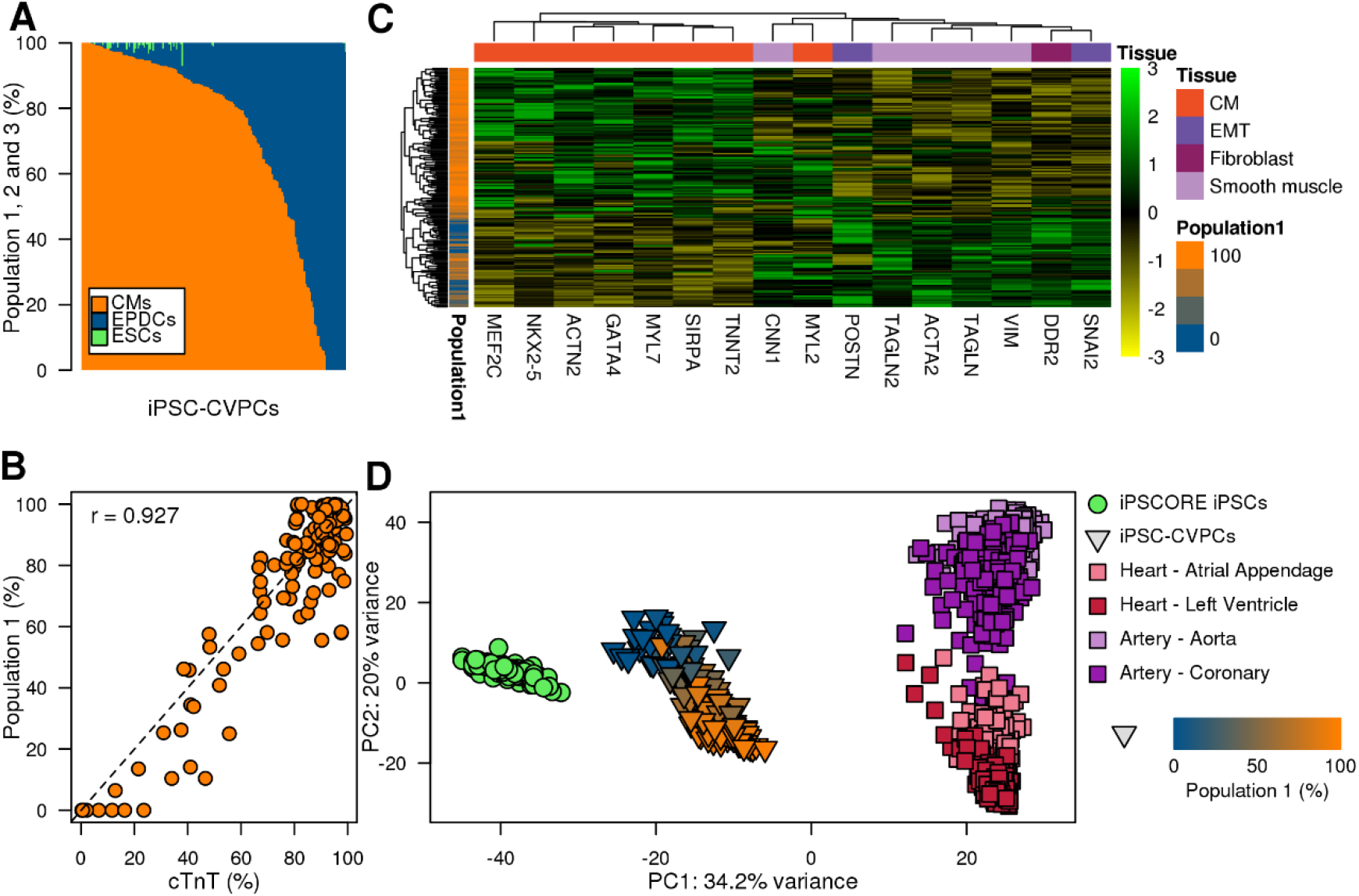
Transcriptomic features of 180 iPSC-CVPC samples. (A) Relative distributions of cell populations estimated using CIBERSORT across 180 iPSC-CVPC samples. (B) Scatterplot showing the correlation between %cTnT (X axis) and the fraction of Population 1 in the iPSC-CVPCs calculated using CIBERSORT (Y axis). (C) Heatmap showing the expression levels of CM and EPDC marker genes (Figure 1K) in 180 iPSC-CVPC samples at D25. (D) PCA on the 1,000 genes with highest variability from 184 iPSC samples, 180 iPSC-CVPC samples (triangles colored according to their % Population 1), and samples from GTEx (squares, left ventricle, right ventricles, coronary artery and aorta).

To characterize the similarities between the iPSC-CVPC transcriptomes and those of adult heart and artery samples, we performed a PCA analysis using the transcriptomes of 184 iPSCORE iPSCs, 180 iPSC-CVPCs, and the 1,072 GTEx samples, including left ventricle, atrial appendage, coronary artery and aorta (Consortium et al., 2017). We found that principal component 1 (PC1) showed that iPSC-CVPCs correspond to an intermediate state between the iPSCs and adult samples, suggesting that the derived CMs and EPDCs are similar to immature cardiac cells (Figure 2D). PC2 divided the samples based on embryonic origins, namely the myocardium (left ventricles and atrial appendages) and epicardium (coronaries and aorta) (Moorman et al., 2003; Perez-Pomares et al., 2016). This analysis shows that derived iPSC-CMs and iPSC-EPDCs lie on different developmental trajectories, with the CMs corresponding to immature myocardium and the EPDCs to immature epicardium.

### iPSC expression signatures impact cardiac fate differentiation

Although all iPSCORE iPSCs have previously been shown to be pluripotent (Panopoulos et al., 2017), we sought to determine if transcriptomic differences existed between the iPSC lines that derived CVPCs containing CMs versus those that gave rise to predominantly EPDCs (Figure 3A). For this analysis, iPSC-CVPCs that completed differentiation (harvested on D25) were defined by the ratio of CM:EPDC estimates from CIBERSORT (estimated %CM: estimated %EPDC), while the 65 iPSC-CVPC differentiations terminated prior to D25 for not forming a beating syncytium were assigned a CM:EPDC ratio of 0:100 (0% CM:100% EPDC). We tested varying CM:EPDC ratios and determined the 30:70 (CM:EPDC) threshold was best for grouping iPSC samples into those that were CM-fated (produced ≥30% CMs), and those that were EPDC-fated (produced >70% EPDCs) (Figures S7-S9, Tables S9, S10; see Methods). We identified 84 differentially expressed genes, 35 of which were overexpressed in the CM-fated iPSC lines and 49 overexpressed in the EPDC-fated iPSCs (Figure 3B,C). These genes have functions associated with three differentiation signatures: 1) Wnt/β-catenin pathway (13 genes); 2) muscle and/or cardiac differentiation (six genes); and 3), EMT and/or mesenchymal tissue development (six genes, Figure 3D, Table S11). We noted that seven borderline significant genes were also involved in one of the three represented signatures, and therefore added them to the final list of differentially expressed genes (Table S11). We investigated the associations between the expression levels of the final list of 91 signature genes in the 184 iPSCs and the fraction of CMs in the resulting iPSC-CVPCs using linear regression, and found significant associations for all genes (Figure 3E, Table S12). These results show that, independently from the 30:70 (CM:EPDC) threshold used in the initial differential expression analysis, the expression levels of these signature genes in the 184 iPSCs were significantly associated with differentiation outcome (e.g. CM-or EPDC-fate).

**Figure 3:**
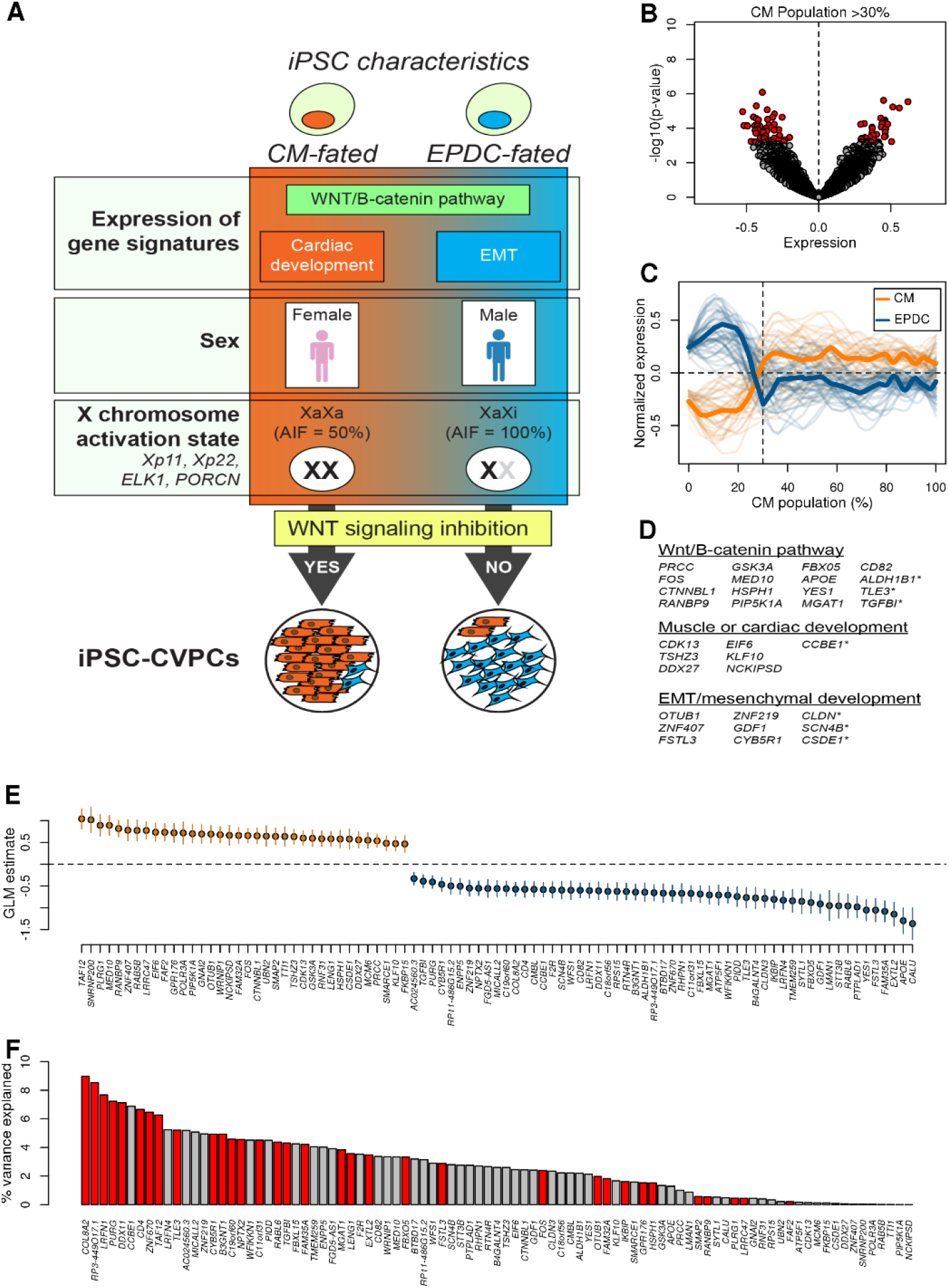
iPSC gene signatures associated with cardiac differentiation fate. (A)Cartoon showing iPSC characteristics that influence their cardiac fate determination, including: 1) the expression levels of 91 genes grouped into three gene signature classes (WNT/B-catenin pathway, cardiac development genes and genes involved in EMT); 2) sex, female iPSCs are more likely to differentiate to CMs than males (see Figure 4); and 3) X chromosome activation state, female iPSCs that have activated both X chromosomes (XaXa) are more likely to differentiate to CMs (see Figure 4). (B) Volcano plot showing mean difference in expression levels for all autosomal genes between CM-fated iPSC lines (their corresponding derived samples have CM population > 30%) and EPDC-fated iPSC lines (X axis) and p-value (Y axis, t-test). A positive difference indicates over-expression in CM-fated iPSCs, whereas a negative difference indicates over-expression in EPDC-fated iPSCs. Significant genes are indicated in red. (C) Expression levels of the 91 signature genes in iPSCs as a function of the % CM population in their corresponding iPSC-CVPC samples. Thick lines represent the average for 36 genes overexpressed in CM-fated iPSCs (orange) and for 55 genes overexpressed in EPDC-fated iPSCs (blue). (D) WNT/β-catenin pathway, muscle/cardiac related, or EMT/mesenchymal development signature genes (those differentially expressed with nominal p-values (p < 0.0015) indicated with an asterisk). (E) GLM estimate (% CM population ~ expression) calculated for each signature gene. Mean and 95% confidence interval are shown. (F) Bar plot showing the percentage of variability in iPSC fate that is explained by each of the 91 signature genes. Bars highlighted in red show the 35 signature genes identified by L1 normalization that independently contributed to variance.

### WNT and differentiation expression signatures contribute to cell fate determination

While the signature genes likely impacted cardiac fate determination, we did not expect each gene to contribute equally. To explore the impact of each gene individually on differentiation outcome, we calculated how much the 91 genes explained the variability underlying iPSC cell fate. To quantify the percent of variance explained by each gene (R^2^), we fit a generalized linear regression model with a logit link function to each gene individually. We found that the percent of variance explained by each individual gene varied over three orders of magnitude (1.73 × 10^−3^ < R^2^ < 8.97%; Figure 3F; Table S12, S13).

We next asked how these signature genes altogether captured variability in differentiation fate. As several of the signature genes had correlated expression levels (Figure S10), to reduce overfitting in the regression analysis, we included an L1 norm penalty (i.e. LASSO regression) and used 10-fold cross validation. We identified 35 genes that independently contributed to variance, and whose expression levels collectively explained more than half of the variability in differentiation outcome across iPSC lines (average R^2^ from the 10-fold cross validation = 0.512; Table S14). Together these data show that, while the proportion of variance explained by each of the signature genes varied widely, altogether they captured approximately half of the total variance underlying differential iPSC fate outcome.

### Inherited genetic variation does not influence differentiation outcome

We investigated if genetic variation associated with the expression of any of the signature genes contributed to the differentiation outcome of iPSCs. We obtained 1,303 genetic variants that had significant associations with the expression levels of at least one of the 91 signature genes from either GTEx (Consortium et al., 2017) or an eQTL analysis using the same collection of iPSCs (DeBoever et al., 2017). We assessed the genotypes of these 1,303 variants in each iPSC line and investigated the association between genotype and the fraction of CMs in the corresponding iPSC-CVPCs. We found that none of these variants were significantly associated with differentiation outcome upon adjustment for multiple testing hypothesis (Storey q-value < 0.1, Table S15, Figure S11). This analysis shows that inherited genetic variants did not contribute to the variance underlying iPSC differentiation outcome captured by the signature genes, indicating that non-genetic factors played a role in their differential expression.

### The role of the X chromosome in cardiac fate differentiation outcome

To understand whether the transcriptomic differences between CM-fated and EPDC-fated iPSCs were associated with alterations in specific pathways or cellular function, we performed a gene set enrichment analysis (GSEA) on 9,808 MSigDB gene sets (Liberzon et al., 2011; Subramanian et al., 2005) using the 15,228 expressed autosomal genes in the 184 iPSCs (see Supplemental text). We identified 22 gene sets that were significantly associated with iPSC cell fate, including enrichment in the CM-fated iPSCs for transcription factor activity, ELK1 targets, and genes in two chromosome X loci (chrXp11 and chrXp22), whereas the EPDC-fated iPSCs were enriched for extracellular matrix (Figure 3A, Figure 4A, Table S16). Notably, the chrXp11 locus encodes both *ELK1* and *PORCN,* whose protein product (Porcupine) is targeted for WNT inhibition during CM differentiation, but not EPDC differentiation (Mo et al., 2013; Wang et al., 2013) (Figure 4B). Supporting the involvement of the X chromosome in differentiation, we observed that sex was strongly associated with differentiation outcome, with female iPSCs being more likely CM-fated and male iPSCs being more likely EPDC-fated (p = 2.57×10^−5^, Z-test, Figure 3A, Figure 4C, Figure S12, Table S17).

**Figure 4:**
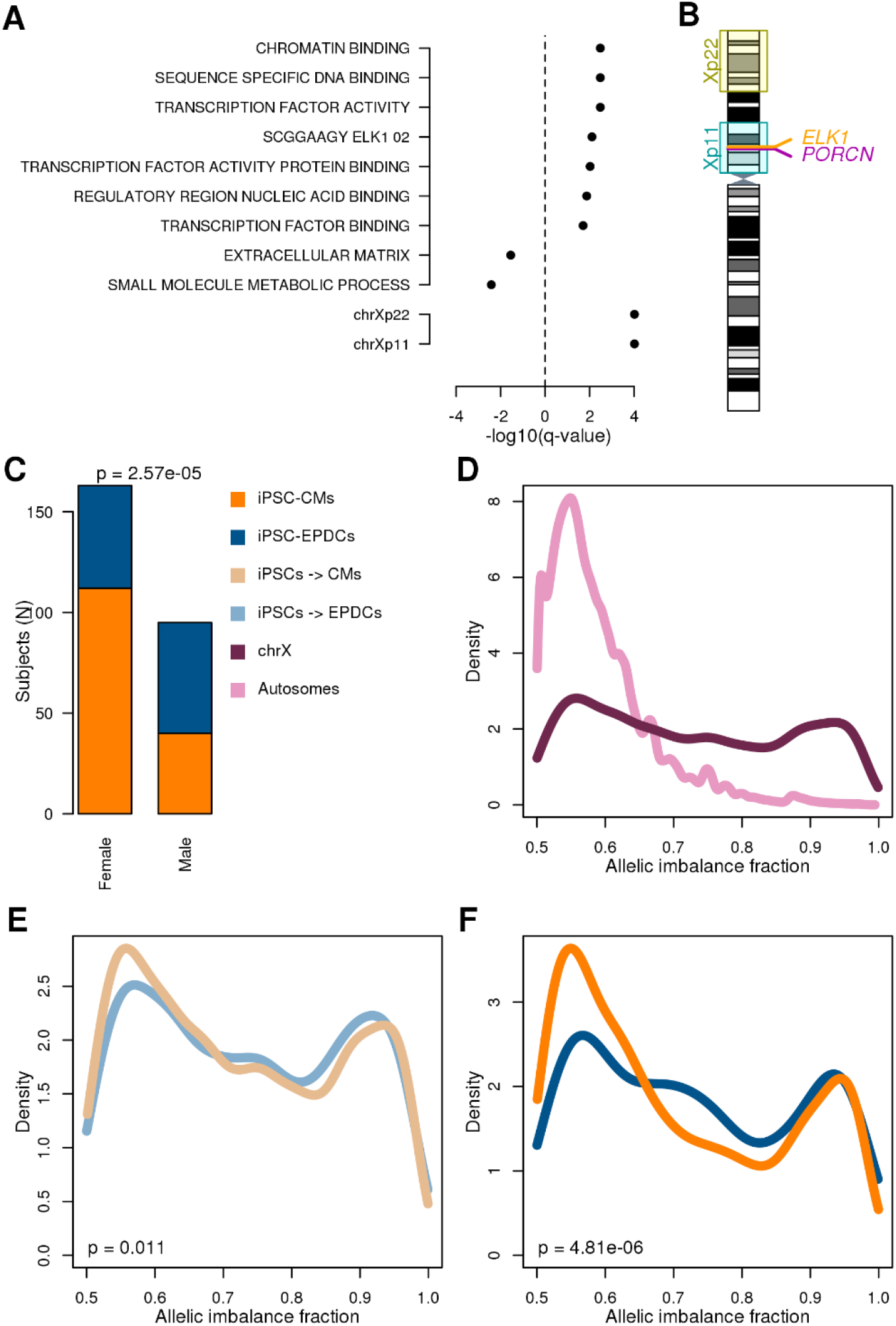
X chromosome gene dosage plays a role in cardiac differentiation fate. (A) GSEA results: For each gene set, -log_10_(q-value) is shown. Positive values correspond to gene sets enriched in CM-fated iPSCs, whereas negative values correspond to EPDC-fated iPSCs. For autosomes all iPSCs were included (top), for the chromosome X only the 113 female iPSCs were analyzed (bottom). (B) Cartoon showing the differentially expressed loci on chromosome X and the position of *ELK1* and *PORCN.* (C) Barplot showing the associations between sex and differentiation outcome (orange: iPSC-CVPC samples with CM fraction > 30%; blue: with EPDC fraction > 70%). P-values were calculated using Z-test (glm function in R). (D-F) Density plots showing the differences in allelic imbalance fraction between (D) autosomal genes (pink) and chrX genes outside of the pseudoautosomal region (maroon) in female iPSCs; (E) chrX genes in female CM-fated (light orange) and EPDC-fated (light blue) iPSCs; and (F) chrX genes in female D25 iPSC-CVPC samples with CM fraction > 30% (orange) and EPDC fraction > 70% (blue). P-values were calculated using Mann Whitney U test.

Given the observation that female iPSCs have a greater potential to differentiate to CMs and that differential expression of chrXp11 genes were associated with differentiation outcome, we asked if variation in X chromosome inactivation (Xi) and activation (Xa) state across female iPSC lines was associated with CM or EPDC-fate. Using RNA-seq data generated from the 113 female iPSCs, we evaluated allele specific effects (ASE) of X chromosome and autosomal genes (Table S18). We defined the strength of ASE for each gene as the fraction of RNA transcripts that were estimated to originate from the allele with higher expression (hereto referred to as “allelic imbalance fraction”, AIF). We observed that AIF in autosomal genes was close to 0.5, indicating that both alleles were equally expressed (Figure 4D), while AIF on the X chromosome in iPSCs tended to be bimodal, with some genes showing monoallelic expression (AIF ~1.0; XaXi) and others showing biallelic expression (AIF ~0.5; XaXa). We observed that AIF was significantly less in the CM-fated female iPSCs compared with the EPDC-fated female iPSCs (p = 0.011, Mann-Whitney U test, Figure 4E) and that this difference in AIF became even more pronounced in the corresponding derived iPSC-CVPC samples (p = 4.81×10^−6^, Mann-Whitney U test, Figure 4F, Figure S13, see Supplemental text). These finding show that differential chromosome XaXi status, as well as altered gene expression in chrXp22 and chrXp11, in iPSCs contribute to differences in cardiac fate differentiation outcome.

### Independent iPSC-CM derivation study validates findings

To assess the generalizability of our findings, we examined an independent collection of 39 iPSCs (Banovich et al., 2018) reprogrammed using an episomal plasmid from Yoruba lymphoblastoid cell lines, and thus were substantially different than the iPSCORE iPSCs (i.e. different reprogramming method, genetic backgrounds, and donor cell types) (Figure 5A, Table S19). Differentiation of these lines using a slightly different protocol resulted in the successful derivation of 15 iPSC-CMs (%cTnT range at D32: 40 to 96.9), whereas 24 were terminated on or before day 10 due to the fact that they did not form a beating syncytium. To examine if the successfully derived Yoruba iPSC-CMs showed the presence of EPDCs, we used RNA-seq data and CIBERSORT to estimate cellular compositions and observed variable relative distributions of CM and EPDC populations (Figure 5B). Consistent with our iPSCORE iPSC-CVPC samples, the estimated CM population fractions were significantly correlated with %cTnT values (r = 0.81, p = 7.94 × 10^−4^, t-test; Figure 5C). Finally, Yoruba iPSC-CMs derived from females tended to have an increased percentage of CMs compared with those derived from males (Figure 5D). These observations show that the Yoruba iPSCs and derived cardiac cells could be used to investigate the generalizability of the associations that we had observed between transcriptomic differences in iPSCs and cardiac fate differentiation outcome.

**Figure 5:**
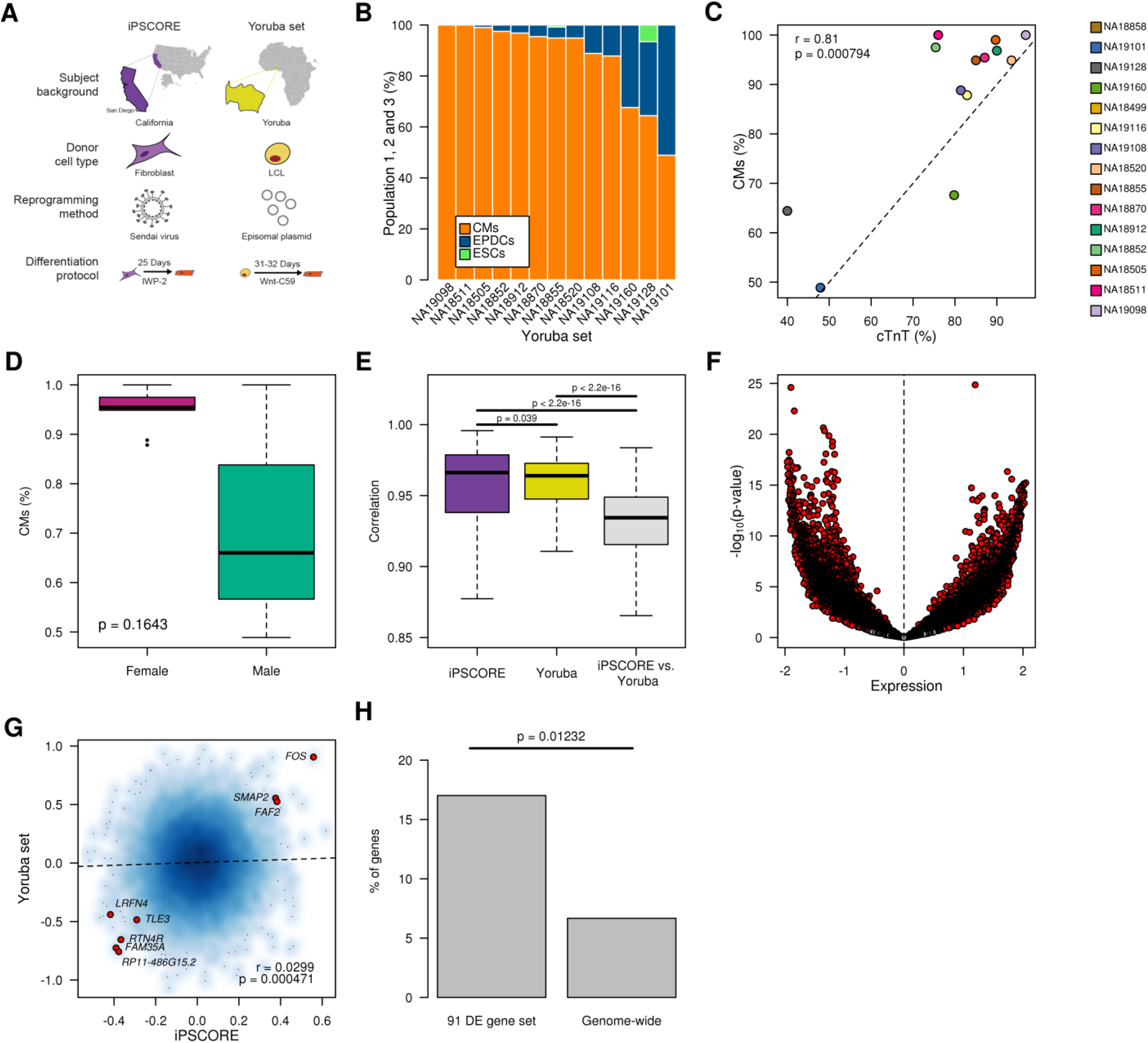
Validation of association between iPSC gene signatures, sex and differentiation outcome. (A) Schematic depicting differences (subject ethnicities, donor cell type, reprogramming method) between the iPSCORE iPSC and Yoruba iPSC as well as differences in the cardiac differentiation protocol (B) Estimated fractions of CMs and EPDCs for 13 Yoruba iPSC-CM samples from RNA-seq using CIBERSORT. (C) Scatterplot showing the correlation between %cTnT (X axis) and the fraction of cells in population 1 (% CMs) calculated using CIBERSORT (Y axis) for 13 Yoruba iPSC-CM samples. (D) Boxplots showing the distribution of estimated fraction of cells in population 1 (% CMs) for 9 female Yoruba iPSC-CM and 4 male Yoruba iPSC-CM. P-value was calculated using Mann-Whitney U test. (E) Box plots showing correlation of gene expression in all 184 iPSCORE iPSCs with RNA-seq (purple), 34 Yoruba iPSCs with RNA-seq used for differentiation (yellow; 14 successful and 20 terminated), and the pairwise comparison of the Yoruba iPSC against the iPSCORE iPSC (grey). (F) Volcano plot showing mean difference in expression levels for all autosomal genes between 14 Yoruba iPSC lines that were successfully differentiated and 125 iPSCORE iPSC lines that differentiated to >30% Population 1 (CMs) (X axis) and p-value (Y axis, t-test). A positive difference between mean expressions indicate iPSCORE-specific over-expression, whereas a negative difference between mean expressions indicate Yoruba-specific over-expression. Significant genes are indicated in red. (G) Smooth color density scatterplot showing gene expression differences between iPSCs with different fates in 184 iPSCORE iPSC (125 CM-fated vs 59 EPDC-fated) (X-axis) to the expression differences between iPSCs with different outcomes in Yoruba iPSC (14 successful vs 20 terminated) (Y-axis). A positive difference indicates shared over-expression of genes between CM-fated iPSC in iPSCORE and successfully differentiated iPSC in the Yoruba set, whereas a negative difference indicates shared over-expression of genes between EPDC-fated iPSC in iPSCORE and terminated iPSC in the Yoruba set. Of the 91 signature genes that were differentially expressed in the iPSCORE iPSCs based on cell fate, eight had nominally significant expression differences in the same direction in the Yoruba iPSC set (shown in red). (H) Barplot showing that the eight iPSCORE differentially expressed genes (panel G) with nominal significant expression differences in the same direction (e.g. over-expressed or down regulated) in the Yoruba iPSCs is greater than random expectation. Of 13,704 genes expressed both in the iPSCORE and Yoruba iPSCs, we obtained 6,909 for which the average normalized expression differences had either the same positive (CM fate/successful differentiation) or negative (EPDC fate/terminated differentiation) direction. The 6,909 genes included 47 of the 91 iPSCORE signature genes. We found that 466 (6.7%) of the 6,909 genes were nominally significant for being differentially expressed between the 14 successful and 20 terminated differentiations in the Yoruba samples, while 8 of the 47 iPSCORE differentially expressed genes (17.0%) had a nominal p < 0.05. This analysis shows that the 91 iPSCORE signature genes are 2.5 times more likely than expected (17.0% vs. 6.7%, p = 0.012, Fisher’s exact test) to be differentially expressed in the Yoruba samples based on cardiac differentiation fate.

As several factors (Figure 5A) were dissimilar between the iPSCORE iPSC and Yoruba iPSC sets, we expected that there would be significant differences between their transcriptional profiles. We initially analyzed how correlated gene expression was: 1) within iPSCORE iPSCs; 2) within Yoruba iPSCs; and 3) between all pairwise comparisons of the iPSCs in these two different collections (Figure 5E). We observed high correlations of gene expression across iPSCs within each collection, however the correlation between samples from different studies was significantly decreased, indicating that the two sets have significant genome-wide gene expression differences. We next examined differential gene expression between the 125 CM-fated iPSCORE iPSC and 14 Yoruba iPSC that successfully differentiated into iPSC-CMs (Figure 5F), and observed that the majority of genes (69.6% with q-value <.10) were significantly differentially expressed between the two iPSC sets. These results show that there are strong batch effects on gene expression between the iPSCORE and Yoruba iPSC lines.

We investigated if, despite the strong batch effects on gene expression between iPSCORE and Yoruba iPSCs, we could detect inherent transcriptional differences impacting cardiac fate determination that were shared between the iPSC sets. Given the relatively small size of the Yoruba study there was insufficient power to detect transcriptional differences between the lines with different differentiation outcomes (Successfully completed versus Terminated). Therefore, for each gene, we compared the mean expression differences between iPSCs with different cardiac fate outcomes in iPSCORE (CM-fate – EPDC-fate) to the expression differences between iPSCs with different differentiation outcomes in the Yoruba set (Successfully completed – Terminated, Figure 5G). We observed a small, but significant correlation (r = 0.0299, p = 4.71 × 10^−4^, t-test) between genes that were differentially expressed in the iPSCORE iPSCs and those that were differentially expressed in the Yoruba iPSCs. Further, we specifically examined the 91 signature genes significantly associated with iPSCORE iPSC cardiac fate outcome and found eight with nominally significant expression differences in the same direction (e.g. overexpressed or downregulated) in the two sets of iPSCs (Figure 5G), which is 2.5 times more than random expectation (p = 0.012, Fisher’s exact test; Figure 5H). These data suggest that the iPSCORE iPSCs and Yoruba iPSCs shared transcriptional differences that impacted cardiac fate differentiation outcome.

## Discussion

The scale of the current study, 258 attempted differentiations of 191 iPSC lines into the cardiac lineage, provided the power to develop a framework to identify non-genetic transcriptional differences that influence cardiac differentiation outcome. Many of the signature genes whose differential expression were associated with differentiation outcome are involved in cardiac development, including the Wnt/β-catenin pathway, muscle differentiation or cardiac-related functions, and the transition of epicardial cells to EPDCs by EMT. We observed that variability across iPSCs on X chromosome gene dosage (XaXa vs XaXi vs XY) played a role in cell fate determination. While iPSCs are known to have only partial XaXa (Anguera et al., 2012; Barakat et al., 2015; DeBoever et al., 2017; Kim et al., 2014; Patel et al., 2017; Sahakyan et al., 2017; Tomoda et al., 2012), we identified two loci (chrXp11 and chrXp22) encoding genes whose expression levels are associated with cardiac lineage fate. The higher expression of chrXp11 genes in CM-fated iPSCs may at least in part be due to fact that *ELK1* and *PORCN* are both encoded in this interval, as the protein product of *PORCN* (Porcupine) is inhibited by IWP-2 during CM differentiation (Mo et al., 2013), but not during EPDC differentiation. Additionally, we found that ELK1 targets are overexpressed in CM-fated iPSCs, which is consistent with previous studies showing that knockdown of *ELK1* in immortalized human bronchial epithelial cells, small airway epithelial cells, and luminal breast cancer cell line (MCF-7) is associated with increased EMT (Desai et al., 2017; Tatler et al., 2016).

Examination of RNA expression in a collection of iPSCs generated from Yoruba individuals showed that female iPSCs tended to be more likely to have a CM-fate and identified shared gene expression differences between the iPSCORE and Yoruba iPSCs associated with cardiac lineage outcome. While some of the non-genetic transcriptomic differences between iPSCs with CM-fates versus those with EPDC-fates observed in iPSCORE may be due to aberrant epigenetic landscapes resulting from the reprogramming method, our observations in the Yoruba iPSCs, which had different genetic backgrounds, donor cell types and reprogramming methods, suggest that our findings will likely be generalizable to other collections of iPSCs.

Overall, our study suggests that expression differences of 91 signature and X chromosome genes result in iPSC lines having differential propensities to respond to WNT inhibition during differentiation, and consequently are fated to produce iPSC-CVPC samples with different proportions of CMs and EPDCs. As iPSCs in both the iPSCORE and Yoruba collections have passed standard quality checks to confirm their pluripotency (Banovich et al., 2018; Panopoulos et al., 2017), these transcriptomic expression differences associated with cardiac lineage outcome are not detected using current quality metrics. In conclusion, our findings suggest that to derive human iPSC lines that respond similarly in differentiation protocols, it may be necessary to improve reprogramming methods such that the transcriptome and X chromosome activation state is fully reset to the naïve state, and incorporate inactivation of one of the X chromosomes in female lines as an early step in differentiation protocols.

## Methods

### iPSCORE subject information

Fibroblasts obtained by skin biopsies from the 181 consented individuals used in this study were recruited as part of the iPSCORE project (Panopoulos et al., 2017). These individuals included seven monozygotic (MZ) twin pairs, members of 32 families (2-10 members/family) and 71 singletons (i.e. not related with any other individual in this study) and were of diverse ancestries: European (118), Asian (27), Hispanic (12), African American (4), Indian (3), Middle Eastern (2) and mix ethnicity (15). The recruitment of these individuals was approved by the Institutional Review Boards of the University of California, San Diego and The Salk Institute (Project no. 110776ZF). Subject descriptions including subject sex, age, family, and ethnicity were collected during recruitment (Table S1). The 181 subjects (108 female and 73 male) used in this study ranged from ages 9 – 88 and include seven monozygotic (MZ) twin pairs, members of 32 families (2-10 members/family) and 71 singletons (i.e. not related with any other individual in this study) and are of various ancestral backgrounds, including 118 European, 27 Asian, 12 Hispanic, four African American, three Indian, two Middle Eastern, and 15 with multiple ethnicities reported. In addition to fibroblast collection for iPSC reprogramming and differentiation, whole blood samples were obtained for whole genome sequencing.

### Whole genome sequencing

As previously described (DeBoever et al., 2017), we generated whole genome sequences from the 181 subjects used for iPSC derivation. Genomic DNA was isolated from whole blood using DNEasy Blood & Tissue Kit (Qiagen) and Qubit quantified. DNA was then sheared using Covaris KE220 instrument and normalized to 1μg, where WGS libraries were prepared using TruSeq Nano DNA HT kit (Illumina) and normalized to 2 – 3.5nM in 6-samples pools. Pooled libraries were clustered and sequenced on the HiSeqX (Illumina; 150 base paired-end) at Human Longevity, Inc. (HLI).

### iPSC derivation

As previously described (Panopoulos et al., 2017), we reprogrammed fibroblast samples from the 181 individuals in this study using non-integrative Cytotune Sendai virus (Life Technologies) (Ban et al., 2011) following the manufacturer’s protocol. The 191 iPSCs used in this study (7 subjects had 2 or more clones each; Table S2) were generated and shown to be pluripotent by analysis of RNA-seq by PluriTest (Muller et al., 2008) and for a subset based on >95% positive double staining for Tra-1-81and SEEA-4 (Panopoulos et al., 2017). They were also shown to have high genomic integrity based on analysis of genomic DNA by SNP arrays (Panopoulos et al., 2017). Cell lysates used for RNA-seq (Tables S5, S6) were collected for 184 of the iPSC samples between passages 12 to 40 (Figure 1A).

### Large-scale derivation of iPSC-CVPC samples

To generate iPSC-derived cardiovascular progenitors (iPSC-CVPCs) we used a small molecule cardiac differentiation protocol (Lian et al., 2013). The 25-day differentiation protocol consisted of five phases (Figure S1A), the optimizations for each step are described in detail below: 1) *expansion:* we developed the ccEstimate algorithm (Figure S2) to automate the detection of 80% confluency for iPSCs in T150 flasks (Figure S1B,C); 2) *differentiation:* we tested whether increasing the dosage of IWP-2 to induce to inhibit the WNT pathway improved differentiation efficiency and found that 7.5 μM at D3 of the differentiation provided in a single dose for 48 hours results in the most efficient differentiation (Figure S1D, E, Table S20); 3)*purification:* since fetal cardiomyocytes use lactate as primary energy source and have a higher capacity for lactate uptake than other cell types (Fisher et al., 1981; Werner and Sicard, 1987), we incorporated lactate metabolic selection for five days to improve iPSC-CVPC purity (Tohyama et al., 2013) (Figure S1F); 4) *recovery:* after metabolic selection, iPSC-CVPCs were maintained in cell culture for five days; and 5) *harvest:* we collected iPSC-CVPCs at D25 for downstream molecular assays and cryopreserved live cells.

The 258 attempted differentiations of the 191 iPSC lines (Table S2) were performed as follows:

#### Expansion of iPSC

One vial of each iPSC line was thawed into mTeSR1 medium containing 10 μM ROCK Inhibitor (Sigma) and plated on one well of a 6-well plate coated overnight with matrigel. During the expansion phase, all iPSC passaging was performed in mTeSR1 medium containing 5 μM ROCK inhibitor, when cells were visually estimated to be at 80% confluency. The iPSCs were passaged using Versene (Lonza) from one well into three wells of a 6-well plate. Next, the iPSCs were passaged using Versene onto three 10 cm dishes at 2.54×10^4^ per cm^2^ density. The iPSCs molonalyer was plated onto three T150 flasks at the density of 3.66 × 10^4^ per cm^2^ using Accutase (Innovative Cell Technologies Inc.). Prior to expansion with Versene, after thaw iPSCs were passaged 1-2 times using Dispase II (20mg/ml; Gibco/Life technologies).

#### Differentiation

At 80% iPSC confluency (measured using ccEstimate, see section below “Estimation of optimal time for initiation of iPSC-CVPCs differentiation using ccEstimate”) cell lysates were collected from 32 lines for RNA-seq data generation, where these iPSC and subsequent generated molecular data are referred to as D0 iPSC (Table S5). After reaching 80% confluency (usually within 4-5 days), differentiation was initiated with the addition of the medium containing RPMI 1960 (gibco-life technologies) with Penicillin – Streptomycin (Gibco/Life Technologies) and B-27 Minus Insulin (Gibco/Life Technologies) (hereafter referred to as RPMI Minus supplemented with 12μM CHIR-99021 (D0). After 24h of exposure to CHIR-99021, medium was changed to RPMI Minus (D1). On D3 medium was changed to 1:1 mix of spent and fresh RPMI Minus supplemented with 7.5μM IWP-2 (Tocris). On D5, after 48h of exposure to IWP-2, the medium was change to RPMI Minus. On D7, medium was changed to RPMI 1960 with with Penicillin – Streptomycin (Gibco/Life Technologies) and B-27 Supplement 50X (hereafter referred to as RPMI Plus) (Gibco/Life Technologies). Between D7 and D13, RPMI Plus medium was changed every 48h.

#### Purification

On D15 the cells were collected from the flask using Accutase and plated onto fresh T150 flasks at confluency 1-1.3 × 10^6^ per cm^2^. On D16, cells were washed with PBS without Ca^2^+ and Mg^2^+ (Gibco/Life Technologies) and medium was changed for RPMI 1960 no glucose (Gibco/Life Technologies) supplemented with Non-Essential Amino Acids (Gibco/Life Technologies), L-Glutamine (Gibco/Life Technologies), Penicillin-Streptomycin 10,000U (Gibco/Life Technologies) and 4mM Sodium L-Lactate (Sigma) in 1M HEPES (Gibco/Life Technologies). Medium supplemented with lactate was changed on D17 and D19.

#### Recovery

On D21 cells were washed with PBS and medium was changed for RPMI Plus. On D23 medium was again changed for RPMI Plus. The first beating cells were usually observed between D7 and D9 and as early as D7 (immediately after the media change) and robust beating was usually observed between D8 and D11. During the lactate selection iPSC-CVPC were beating robustly less than 16 hours after reseeding. For all successfully derived iPSC-CVPCs on D25, total-cell lysate material was collected and frozen for downstream RNA-seq assays.

#### Harvest

On D25 cells were collected using Accuase and processed for the following molecular material for downstream assays: 1) cell lysates (RNA-Seq); 2) permeabilized cells (ATAC-Seq); 3) live frozen cells (scRNA-seq); 4) cross-linked cells (ChIP-Seq, median number of vials/iPSC line = 3; ~1.0 × 10^7^ cells/vial), and 5) dry cell pellets (methylation and protein). RNA-seq was generated from 180 iPSC-CVPC differentiations (149 lines from 139 subjects) that successfully reached D25 (Table S5).

### Estimation of optimal time for initiation of iPSC-CVPCs differentiation using ccEstimate

We developed an automatic pipeline that analyzes images of monolayer-grown cells and determines their confluence (Figures S1C, S2). <u>C</u>ell <u>c</u>onfluency <u>estimates</u> (ccEstimate) are performed by first dividing each T150 flask into 10 sections (Figure S1C) and acquiring images for each section every 24 hours after cells are plated as a monolayer. The final image is acquired immediately after treatment with CHIR, which occurs when their confluence is at least 80% (Day 0). The time required for cells to reach 80% confluence is estimated on the basis of the confluence curve derived for each section in each flask. To digitally measure iPSC confluency, ccEstimate performs image analysis using the EBImage package in R (Pau et al., 2010). Images are read using the readImage function. As lighting may be different between the center and the border of an image, only the central part of the image is retained. To separate cells from the background and calculate confluence (i.e. the fraction of the surface of the flask that is covered by cells) the following operations are performed (Figure S2):

1. The image is transformed to monochromatic by determining the intensity of each pixel as the average of the intensities of the red, green and blue channels.
2. Edges are sharpened using high-pass filter. The matrix used for this filter is 15×15 with values −1 on the diagonals and +28 in the center.
3. Contrasts are enhanced by multiplying the pixel intensities by 2.
4. Mean and standard deviation of the pixel intensities are calculated. The image is transformed from monochromatic to binary by setting all pixels with intensity more than two standard deviations higher than the mean to white (intensity = 1) and all other pixels to black (intensity = 0).
5. The resulting binary image is dilated using a disc-shaped structuring element with diameter 5 pixels.
6. 1,000 50×50 pixels sub-images are randomly selected. For each sub-image, the number of white pixels is calculated. Confluence is estimated as the fraction of the randomly selected sub-images with at least 50% of white pixels.

Confluency measurement data is collected for at least the first three days after plating as monolayer to train a generalized linear model (GLM) using the function glm in R to estimate when cells must be treated with CHIR. Estimation is performed separately for each flask section and CHIR is added to all three flasks associated to a given line when at least 75% of sections have confluence 80% (Figure S1C).

### Optimization of IWP-2 concentration by visual estimation of iPSC-CVPCs structure and beating quality

To optimize the IWP-2 concentration, one iPSC line (2_3) was differentiated under four different IWP-2 conditions (Figure S1D, E): 1) 5μM IWP-2 added on D3, 2) 7.5μM IWP-2 added on D3, 3) 5μM IWP-2 added on D3 and D4, or 4) 7.5μM IWP-2 added on D3 and D4. In all four conditions cells were exposed to IWP-2 for 48 hours. At D15 of differentiation, the quality of generated iPSC-CVPC structures and beating were estimated by visual evaluation using two metrics that we established in the lab: 1) structure score; and 2) beat score. Both structure score and beat score were evaluated at 10 spots on each 150T flask that had also been used for digital measurement of cell confluency (Table S20). Structure score and beat score had 4-point scales where 0 was the lowest and 3 was the highest grade. For structure score 0 = less than 10% of cells were cardiomyocyte-like with thick structures; 1 = 10-25% of cells were cardiomyocyte-like with thick structures; 2 = over 50% of cells were cardiomyocyte-like with thick structures; 3 = over 90% of cells were cardiomyocyte-like with thick structures.

For beat score 0 = less than 10% of cells were cardiomyocyte-like beating robustly as a sheet; 1= 10-25% of cells were cardiomyocyte-like beating robustly; 2 = over 50 of cells were cardiomyocyte-like beating robustly; 3 = over 90% of cells were cardiomyocyte-like beating robustly. In cases of uncertainty or intermediate results, cells were assigned a lower grade. Grade 3 was assigned only for the iPSCs with thick, robustly beating sheets of cells.

### Comparison of lactate and glucose treated iPSC-CVPCs

To examine the effects of lactate purification, three iPSC-CVPC lines derived from unrelated individuals (2_3, 8_2, and 3_2) were differentiated to D15 (Figure S1F). At D16, medium supplemented with either 4mM Sodium L-Lactate (Sigma) or 2mg/mL D-glucose (Gibco/Life Technologies). Medium was changed on D17 and D19. On D21 cells were washed with PBS and medium was changed for RPMI Plus. Lactate and glucose treated cells were harvested on D25.

### Flow cytometry

On D25 of differentiation, 5×10^5^ iPSC-CVPCs were permeabilized and blocked in 0.5% BSA, 0.2% TX-100 and 5% goat serum in PBS for 30 minutes at room temperature. Cells were stained with Troponin T, Cardiac Isoform Ab-1, Mouse Monoclonal Antibody (Thermo Scientific, MS-295-P0) at 4°C for 45 minutes, followed by Alexa Fluor 488 secondary antibody (Life Technologies, A11001). Stained cells were acquired using BD FACSCanto II system (BD Biosciences) and analyzed using FlowJo V10.2.

### Immunofluorescence analysis of iPSC-CVPCs

Immunofluorescence (IF) was assessed in 5 iPSC-CVPC lines (13_1, 14_2, 29_1, 2_1, and 42_1). Cells for IF were obtained by thawing live frozen iPSC-CVPC harvested on D25 and plating them directly on 0.1% gelatin-coated glass-bottom plates for five days (D30). Cells were then fixed using 4% paraformaldehyde (PFA) in PBS or 20 min at room temperature (RT). Fixed cells were permeabilized for 8 min at RT with 0.1% Triton X-100 in PBS, blocked in 5% bovine serum albumin for 30 min at RT and incubated overnight at 4°C with a primary antibody. Cells were incubated with rabbit polyclonal anti-connexin 43 (Cx43) antibody (Invitrogen, 710700) and with mouse monoclonal anti-sarcomeric alpha-actinin antibody (Sigma, A7811), or with rabbit polyclonal anti-MLC2V (Proteintech, 10906-1-AP) and/or mouse monoclonal anti-MLC2A (Synaptic Systems, 311011). All antibodies are described in Table S4.

After overnight incubation cells were washed three times with PBS and incubated with appropriate secondary antibodies: donkey anti-rabbit Alexa Fluor 488 (Invitrogen, A-21206) and goat anti-mouse Alexa Fluor 568 (Invitrogen, A-11004) secondary antibodies for 45 mins at RT. Cells were washed three times with PBS and nuclei were counterstained with DAPI and mounted. Slides were imaged using Olympus FluoView FV1000 confocal microscope at UCSD Microscopy Core.

### Generation of RNA-seq data

For gene expression profiling of iPSCs, we used RNA-seq data from 184 samples that we previously published (DeBoever et al., 2017). We generated additional 180 RNA-seq samples from iPSC-CVPC samples at D25 differentiation (Table S6). All RNA-seq samples were generated and analyzed using the same pipeline (DeBoever et al., 2017). Briefly, we isolated total RNA from total-cell lysates using the Quick-RNA™ MiniPrep Kit (Zymo Research) from frozen total-cell lysate, including on-column DNAse treatment steps and eluted in 48 μl RNAse-free water. RNA elutions were run on a Bioanalyzer (Agilent) to determine integrity and all samples had RNA integrity number (RIN) values greater than 9. Illumina Truseq Stranded mRNA libraries were prepared and sequenced on HiSeq4000, to an average of 28 M 125 bp paired-end reads per sample. RNA-Seq reads were aligned using STAR60 with a splice junction database built from the Gencode v19 gene annotation. RNA-Seq data with percent uniquely mapped reads greater than 70% and percent duplication less than 50% were considered to be good quality. Transcript and gene-based expression values were quantified using the RSEM package (1.2.20) (Li and Dewey, 2011) and normalized to transcript per million bp (TPM).

### Generation of scRNA-seq data

#### Generation

For eight iPSC-CVPCs sample and one H9 ESC line, single cells were captured using the 10× Chromium controller (10× Genomics) according to the manufacturer’s specifications and manual (Manual CG00052, Rev C). Cells for each sample were loaded on the individual lane of a Chromium Single Cell A Chip. Libraries were generated using Chromium Single Cell 3’ Library Gel Bead Kit v2 (10× Genomics) following manufactures manual. Libraries were sequenced using a custom program (26-8-98 Pair End) on HiSeq 4000. Each library was sequenced on an individual lane. In total we captured 36,839 cells. We retrieved FASTQ files and used CellRanger V2.1 (https://support.10xgenomics.com/) with default parameters using Gencode V19 gene annotation to generate single-cell gene counts for each individual sample.

#### Processing

To combine the scRNA-seq from each individual sample, we used *cellranger aggr* and obtained a total of 36,839 cells from 8 iPSC-CVPCs and 1 ESC sample. We removed 1,934 cells because they were not in G0 phase, as they expressed the proliferation marker MKI67 (Scholzen and Gerdes, 2000) at high levels (UMI > 2, Figure S4A-D). We also removed doublets (i.e. sequenced droplets containing more than one cell)(Kang et al., 2018) by visual inspection of the t-SNE plots (Figure S4). There were 34,905 cells remaining after proliferating cells and doublets were removed. K-means clustering was performed on the 34,905 cells using k values 3, 4, and 9 (Figure S4E-G). k = 3 was determined to be the most suitable value, as visual inspection of the principal component analysis showed 3 distinct clusters (Figure S4E).

#### Differential expression

Differential expression across the three scRNA-seq clusters was performed by comparing the distribution of unique molecular identifiers (UMI) for a given gene from all the cells specific to one cluster (k-means; k = 3) with all the cells specific to the other two clusters using edgeR asymptotic beta test (Robinson and Smyth, 2008) (Table S8). Differentially genes that had a total UMI ≥ 1 and FDR < 0.05 were considered to be significantly overexpressed in a given cluster. For visualization of gene expression in the t-SNE plots, transcript levels for each gene were normalized using the *calcNormFactors* function in edgeR (Robinson et al., 2010).

### CIBERSORT

The expression levels of the top 50 genes overexpressed in each of the three cell populations (total 150 genes), with nominal p-value < 1.0 × 10^−13^ and mean UMI > 1 (Table S7), were used as input for CIBERSORT (Newman et al., 2015) to calculate the relative distribution of the three cell populations for all the 180 iPSC-CVPC samples at D25. CIBERSORT (https://cibersort.stanford.edu/) was run with default parameters using the TPM values for the 150 genes in all 180 iPSC-CVPC samples.

### Characterizing transcriptional similarities of iPSCs, iPSC-CVPCs and GTEx adult tissues by principal component analysis

We performed principal component analysis (PCA) on RNA-seq using R prcomp function on 184 iPSCs, 180 iPSC-CVPCs and 1,072 RNA-seq samples from GTEx, including 303 left ventricle samples, 297 atrial appendage samples, 173 coronary artery samples and 299 aorta samples.

### Identifying differentially expressed genes between iPSCs with different cardiac fates

For differential expression analysis, for each line that had more than one iPSC-CVPC differentiation, we used the sample with the highest Population 1 fraction.

To calculate differential expression between CM-fated iPSCs and EPDC-fated iPSCs, we first retained all genes with TPM ≥ 2 in at least 10 samples and then transformed the RNA-seq TPM data to standard normal distributions by quantile normalization using the function normalize.quantiles from R package preprocessCore (Bolstad et al., 2003). Quantile normalized expression levels were then corrected for the first 10 factors calculated by PEER (Stegle et al., 2012).

To obtain a cutoff, we used the RNA-seq data to conduct a series of differential expression analyses on 15,228 autosomal genes in the 184 iPSC lines (147 completed and 37 terminated) with RNA-seq data considering the ratio of population frequencies in the corresponding derived iPSC-CVPCs (0:100, 10:90, 20:80, 30:70, 40:60, 50:50, 60:40, 70:30, 80:20 and 90:10) (Table S9, Figure S7); for example, for the 90:10 ratio we compared gene expression in the 184 iPSCs that differentiated into iPSC-CVPCs with >= 90% Population 1 to iPSCs that differentiated into iPSC-CVPCs with less than 90% Population 1. Across the thresholds, the top differentially expressed autosomal genes (t-test) were always the same (Table S9); however, the 30:70 (Population 1:Population 2) ratio resulted in the highest number of differentially expressed genes (93 genes with Storey q-value < 0.1, t-test, Figures 4A and S7, Table S10). Thus, we grouped the 184 iPSC lines into: 1) those that have CM fates, i.e. produced iPSC-CVPC with >= 30% Population 1 (125 lines), and 2) those that have EPDC fates, i. e. produced iPSC-CVPC with > 70% Population 2 (22 lines differentiated to D25 and 37 terminated lines). To remove any biases resulting from the fact that the ratio of male to female iPSCs was 71:113, we filtered the 93 genes removing those that were significantly (q-value < 0.1, t-test) differentially expressed between female and male iPSCs, resulting in a set of 84 genes, which is substantially greater than random expectation (Figure S8).

To determine if the number of significantly differentially expressed genes was higher than expected by chance, we shuffled the assignments of the 184 iPSC RNA-seq samples to differentiation fate (125 CM and 59 EPDC) 100 times. For each shuffle, we performed differential expression analysis and obtained the number of genes that were significantly differentially expressed. In Figure S9 we show a QQ plot that demonstrates that the observed p-value distribution was substantially different than random expectation.

### Contribution of 91 signature genes in iPSCs to determination of cardiac fate

#### Individual contributions

For each of the 91 signature genes, we built a generalized linear model (GLM) with the expression of the gene as input and the differentiation outcome (e.g. % Population 1) as output using the LinearRegression function from sklearn. To model the continuous property of the % Population 1 distributions, but maintain their boundary from 0-100, we used a logit link function to transform measurements of % cardiomyocyte to ln(OR) of % Population 1, calculated as ln(% Population 1/ (1 – % Population 1) and capped the percentages at 0.99 and 0.01 to avoid infinite or undefined odds ratios. For each gene, the percent of variance explained is defined as the model’s R^2^.

#### Cumulative impact

To understand the cumulative contribution of all 91 signature genes on cardiac differentiation fate, we built a generalized linear model (GLM) with an L1 norm penalty (ie LASSO) using the expression of all 91 genes as input and the differentiation outcome (e.g. % Population 1) as output using the LassoLarsCV function from sklearn. To model the continuous property of the % Population 1 distributions, but maintain their boundary from 0-100, we used a logit link function to transform measurements of % cardiomyocyte to ln(OR) of % Population 1, calculated as ln(% Population 1/ (1 – % Population 1) and capped the percentages at 0.99 and 0.01 to avoid infinite or undefined odds ratios. To avoid overfitting the model, we used10-fold cross validation implemented in sci-kit learn v0.19.1 with 10,000 max iterations (Pedregosa et al., 2011).

### Detecting associations between genetic variation and differentiation outcome

eQTL data was retrieved for all GTEx V.7 tissues(Consortium et al., 2017) and for iPSCs (DeBoever et al., 2017). We obtained 1,795 variants that were eQTLs for any of the 91 signature genes, of which 1,303 had minor allelic frequency (MAF) > 1% in the 181 individuals from whom iPSCs were derived. Genotypes were obtained for each SNP in all individuals using *bcftools view* (Li, 2011) and multiallelic variants were decomposed to monoallelic using *vt decompose* (Tan et al., 2015). Linear regression was used to calculate the associations between the genotype of each variant and differentiation outcome (% CM population in the iPSC-CVPCs).

### Gene set enrichment analysis using the MSigDB collection

We performed gene set enrichment analysis (GSEA) using the R *gage* package (V 2.20.1) (Luo et al., 2009) on all MSigDB gene sets (Liberzon et al., 2011; Subramanian et al., 2005) from the 8 collections, including Hallmark gene sets (H), positional gene sets (C1), curated gene sets (C2), motif gene sets (C3), computational gene sets (C4), Gene Ontology (GO, C5), oncogenic signatures (C6), and immunologic signatures (C7). FDR correction was performed independently for each collection (see Supplemental text). The normalized mean expression difference between iPSCs that differentiated to CMs and iPSCs that differentiated to EPDCs (Table S10) was used as input for GSEA. Gene lists that were significant after multiple testing correction (FDR p < 0.05) were considered significant.

### Associations between iPSC and subject features and differentiation outcome

A generalized linear model (GLM) was built in R using age, sex, ethnicity, age, and passage of the iPSCs at D0 of differentiation as input and differentiation outcome as output (0 = EPDCs; and 1 = CMs). The model was built using the function glm(outcome ~ age + sex + ethnicity + passage, family=binomial(link=‘logit’)).

### Identifying X chromosome inactivation in female iPSCs and iPSC-CVPCs

To analyze X chromosome inactivation, we used 113 female iPSCs, of which 87 where CM-fated and 26 were EPDC-fated (see Supplemental text). To call allele specific effects (ASE) in RNA-Seq from iPSC and iPSC-CVPCs, we used the method previously described in DeBoever *et al.* (DeBoever et al., 2017). Genes lying in X chromosome pseudoautosomal (PAR) regions (PAR1: 60001-2699520, PAR2: 154931044 – 155260560) were removed from the analysis. We defined the strength of ASE for each gene as the fraction of RNA transcripts that were estimated to originate from the allele with higher expression (referred to as allelic imbalance fraction, AIF).

### Validation of findings in Yoruba iPSC set

#### Generation of iPSCs

The Yoruba iPSCs in the Banovich *et al.* study (Banovich et al., 2018) were generated from lymphoblastoid cell lines (LCLs) using an episomal reprogramming strategy. Briefly, this included transfecting LCLs with the episomal plasmids and then culturing for seven days in hESC media (DMEM/F12 supplemented with 20% KOSR, 0.1 mM NEAA, 2mM GlutaMAX, 1% Pen/Strep, 0.1# 2-Mercaptoethanol, 25ng/μl of bFGF, and 0.5mM NaB). On day eight, the transfected cells were plated in a 6-well plates. After four days, NaB was removed from the hESC media. Colonies were observed within 21 days and passaging continued for an additional 10 weeks (1 passage / week), where cells were collected for cryopreservation. Material collected for RNA-seq of the iPSC were collected after an additional minimum of three passages.

#### Differentiation protocol

The Yoruba iPSC-CM derivation (Banovich et al., 2018) was performed using a small molecular method similar to iPSCORE iPSC differentiation protocol (see above: Large-scale iPSC-CVPC deviation). Briefly, 39 iPSCs were expanded until 70-100% confluency (three to five days). On D0, differentiation was initiated by the supplementation of media with 12μM of GSK3 inhibitor CHIR-99021 for WNT pathway activation. On D3 of differentiation, 2μM of Wnt-C59 was added (PORCN inhibitor). On D5 of differentiation, Wnt-C59 was removed from culturing media and differentiating cells were grown with regular media exchanges from D5 to D14. On D14, D16, and D18 cultures were exposed to 5mM Sodium L-lactate for cardiomyocyte purification. On D20-D25, differentiating cells were exposed to 1.7 mg/mL galactose daily to force aerobic metabolism and thus aid in cardiomyocyte maturation. On D25-D27, cells were incubated at physiological oxygen levels (10%). On D27 cells were electrically stimulated with 6.6 V/cm, 2ms and 1Hz for further aid in cardiomyocyte maturation. Finally, iPSC-CMs were harvested on D31 or D32. Purity of iPSC-CM Yoruba lines were measured by cTnT marker and flow cytometry. Out of the 39 iPSCs for which differentiation was attempted, 15 lines successfully generated iPSC-CMs and 24 were terminated on or before day 10 due to the fact that they did not form a beating syncytium (Table S19).

#### RNA-seq

We downloaded RNA-seq for 34 iPSC (five iPSCs did not have RNA-seq) and 13 iPSC-CM samples (two iPSC-CMs did not have RNA-seq) from Gene Expression Omnibus (GEO; GSE89895) (Banovich et al., 2018). Briefly, these Yoruba RNA-seq data were generated from Illumina TrueSeq prepared libraries and sequenced at 50bp single-end reads on an Illumina 2500. As iPSCORE RNA-seq was 125 bp paired-end reads, for comparative analyses, we trimmed all iPSCORE iPSC and iPSC-CM data to 50 bp and treated the paired-end reads as single-end reads. Both iPSCORE and Yoruba 50 bp RNA-seq was then processed as described above (Methods: Generation of RNA-seq data). Briefly, RNA-seq was aligned using STAR60, then gene expression was quantified using the RSEM package and normalized to TPM.

#### Estimation of cellular composition

The RNA-seq for the 13 Yoruba iPSC-CMs were analyzed using CIBERSORT similar to the iPSCORE samples (see CIBERSORT section above). Briefly, the TPM values of the 150 overexpressed genes (50 from each of the three single cell populations; Table S8) in the 13 Yoruba iPSC-CM were used as input to CIBERSORT to calculate the relative distribution of the three populations.

## Data and software availability

The accession numbers for the RNA-seq data, scRNA-seq, and whole-genome sequence genotypes reported in this paper are dbGaP: phs00924 and phs001325. The 191 iPSC lines are available through WiCell Research Institute: https://www.wicell.org/; NHLBI Next Gen Collection.

## Acknowledgements

This work was supported in part by a California Institute for Regenerative Medicine (CIRM) grant GC1R-06673 and NIH grants HG008118-01, HL107442-05, DK105541-03, and DK112155-01. RNA-seq were performed at the UCSD IGM Genomics Center with support from NIH grant P30CA023100. M.K.R.D. was supported by the National Library of Medicine Training Grant T15LM011271. W.W.G. was supported by the National Heart, Lung, And Blood Institute of the National Institutes of Health under Award Number F31HL142151.

## Author information

K.A.F., A.D.C., M.K.R.D., and M.D. conceived the study. A.D.C. performed the iPSC-CVPC differentiations. A.D.C. and P.B. generated the molecular data. S.H. generated the IF images. M.C.W. generated iPSC-CMs from Yoruba iPSCs. F.S and A.D.C generated scRNA-seq data. M.K.R.D., H.M.., E.N.S, and M.D. performed data processing and computational analyses. M.K.R.D., M.D., and W.W.G. performed gene expression analysis. M.P., E.A., and K.A.F. oversaw the study. M.K.R.D., M.D., and K.A.F. prepared the manuscript.

## References

Anguera, M.C., Sadreyev, R., Zhang, Z., Szanto, A., Payer, B., Sheridan, S.D., Kwok, S., Haggarty, S.J., Sur, M., Alvarez, J., et al. (2012). Molecular signatures of human induced pluripotent stem cells highlight sex differences and cancer genes. Cell stem cell 11, 75–90.

Ban, H., Nishishita, N., Fusaki, N., Tabata, T., Saeki, K., Shikamura, M., Takada, N., Inoue, M., Hasegawa, M., Kawamata, S., et al. (2011). Efficient generation of transgene-free human induced pluripotent stem cells (iPSCs) by temperature-sensitive Sendai virus vectors. Proceedings of the National Academy of Sciences of the United States of America 108, 14234–14239.

Banovich, N.E., Li, Y.I., Raj, A., Ward, M.C., Greenside, P., Calderon, D., Tung, P.Y., Burnett, J.E., Myrthil, M., Thomas, S.M., et al. (2018). Impact of regulatory variation across human iPSCs and differentiated cells. Genome research 28, 122–131.

Bao, X., Lian, X., Hacker, T.A., Schmuck, E.G., Qian, T., Bhute, V.J., Han, T., Shi, M., Drowley, L., Plowright, A., et al. (2016). Long-term self-renewing human epicardial cells generated from pluripotent stem cells under defined xeno-free conditions. Nat Biomed Eng 1.

Barakat, T.S., Ghazvini, M., de Hoon, B., Li, T., Eussen, B., Douben, H., van der Linden, R., van der Stap, N., Boter, M., Laven, J.S., et al. (2015). Stable X chromosome reactivation in female human induced pluripotent stem cells. Stem Cell Reports 4, 199–208.

Bolstad, B.M., Irizarry, R.A., Astrand, M., and Speed, T.P. (2003). A comparison of normalization methods for high density oligonucleotide array data based on variance and bias. Bioinformatics 19, 185–193.

Burridge, P.W., Matsa, E., Shukla, P., Lin, Z.C., Churko, J.M., Ebert, A.D., Lan, F., Diecke, S., Huber, B., Mordwinkin, N.M., et al. (2014). Chemically defined generation of human cardiomyocytes. Nature methods 11, 855–860.

Carcamo-Orive, I., Hoffman, G.E., Cundiff, P., Beckmann, N.D., D’Souza, S.L., Knowles, J.W., Patel, A., Papatsenko, D., Abbasi, F., Reaven, G.M., et al. (2017). Analysis of Transcriptional Variability in a Large Human iPSC Library Reveals Genetic and Non-genetic Determinants of Heterogeneity. Cell stem cell 20, 518–532 e519.

Consortium, G.T., Laboratory, D.A., Coordinating Center-Analysis Working, G., Statistical Methods groups-Analysis Working, G., Enhancing, G.g., Fund, N.I.H.C., Nih/Nci, Nih/Nhgri, Nih/Nimh, Nih/Nida, et al. (2017). Genetic effects on gene expression across human tissues. Nature 550, 204–213.

DeBoever, C., Li, H., Jakubosky, D., Benaglio, P., Reyna, J., Olson, K.M., Huang, H., Biggs, W., Sandoval, E., D’Antonio, M., et al. (2017). Large-Scale Profiling Reveals the Influence of Genetic Variation on Gene Expression in Human Induced Pluripotent Stem Cells. Cell stem cell 20, 533–546 e537.

Desai, K., Aiyappa, R., Prabhu, J.S., Nair, M.G., Lawrence, P.V., Korlimarla, A., Ce, A., Alexander, A., Kaluve, R.S., Manjunath, S., et al. (2017). HR+HER2-breast cancers with growth factor receptor-mediated EMT have a poor prognosis and lapatinib downregulates EMT in MCF-7 cells. Tumour Biol 39, 1010428317695028.

Dubois, N.C., Craft, A.M., Sharma, P., Elliott, D.A., Stanley, E.G., Elefanty, A.G., Gramolini, A., and Keller, G. (2011). SIRPA is a specific cell-surface marker for isolating cardiomyocytes derived from human pluripotent stem cells. Nature biotechnology 29, 1011–1018.

Fisher, D.J., Heymann, M.A., and Rudolph, A.M. (1981). Myocardial consumption of oxygen and carbohydrates in newborn sheep. Pediatric research 15, 843–846.

Friedman, C.E., Nguyen, Q., Lukowski, S.W., Helfer, A., Chiu, H.S., Miklas, J., Levy, S., Suo, S., Han, J.J., Osteil, P., et al. (2018). Single-Cell Transcriptomic Analysis of Cardiac Differentiation from Human PSCs Reveals HOPX-Dependent Cardiomyocyte Maturation. Cell stem cell 23, 586–598 e588.

Guzzo, R.M., Gibson, J., Xu, R.H., Lee, F.Y., and Drissi, H. (2013). Efficient differentiation of human iPSC-derived mesenchymal stem cells to chondroprogenitor cells. J Cell Biochem 114, 480–490.

Hartman, M.E., Dai, D.F., and Laflamme, M.A. (2016). Human pluripotent stem cells: Prospects and challenges as a source of cardiomyocytes for in vitro modeling and cell-based cardiac repair. Adv Drug Deliv Rev 96, 3–17.

Iyer, D., Gambardella, L., Bernard, W.G., Serrano, F., Mascetti, V.L., Pedersen, R.A., Talasila, A., and Sinha, S. (2015). Robust derivation of epicardium and its differentiated smooth muscle cell progeny from human pluripotent stem cells. Development 142, 1528–1541.

Kang, H.M., Subramaniam, M., Targ, S., Nguyen, M., Maliskova, L., McCarthy, E., Wan, E., Wong, S., Byrnes, L., Lanata, C.M., et al. (2018). Multiplexed droplet single-cell RNA-sequencing using natural genetic variation. Nature biotechnology 36, 89–94.

Kattman, S.J., Witty, A.D., Gagliardi, M., Dubois, N.C., Niapour, M., Hotta, A., Ellis, J., and Keller, G. (2011). Stage-specific optimization of activin/nodal and BMP signaling promotes cardiac differentiation of mouse and human pluripotent stem cell lines. Cell stem cell 8, 228–240.

Kilpinen, H., Goncalves, A., Leha, A., Afzal, V., Alasoo, K., Ashford, S., Bala, S., Bensaddek, D., Casale, F.P., Culley, O.J., et al. (2017). Common genetic variation drives molecular heterogeneity in human iPSCs. Nature 546, 370–375.

Kim, K.Y., Hysolli, E., Tanaka, Y., Wang, B., Jung, Y.W., Pan, X., Weissman, S.M., and Park, I.H. (2014). X Chromosome of female cells shows dynamic changes in status during human somatic cell reprogramming. Stem Cell Reports 2, 896–909.

Li, B., and Dewey, C.N. (2011). RSEM: accurate transcript quantification from RNA-Seq data with or without a reference genome. BMC Bioinformatics 12, 323.

Li, H. (2011). A statistical framework for SNP calling, mutation discovery, association mapping and population genetical parameter estimation from sequencing data. Bioinformatics 27, 2987–2993.

Lian, X., Bao, X., Zilberter, M., Westman, M., Fisahn, A., Hsiao, C., Hazeltine, L.B., Dunn, K.K., Kamp, T.J., and Palecek, S.P. (2015). Chemically defined, albumin-free human cardiomyocyte generation. Nature methods 12, 595–596.

Lian, X., Zhang, J., Azarin, S.M., Zhu, K., Hazeltine, L.B., Bao, X., Hsiao, C., Kamp, T.J., and Palecek, S.P. (2013). Directed cardiomyocyte differentiation from human pluripotent stem cells by modulating Wnt/beta-catenin signaling under fully defined conditions. Nature protocols 8, 162–175.

Liberzon, A., Subramanian, A., Pinchback, R., Thorvaldsdottir, H., Tamayo, P., and Mesirov, J.P. (2011). Molecular signatures database (MSigDB) 3.0. Bioinformatics 27, 1739–1740.

Luo, W., Friedman, M.S., Shedden, K., Hankenson, K.D., and Woolf, P.J. (2009). GAGE: generally applicable gene set enrichment for pathway analysis. BMC Bioinformatics 10, 161.

Mo, M.L., Li, M.R., Chen, Z., Liu, X.W., Sheng, Q., and Zhou, H.M. (2013). Inhibition of the Wnt palmitoyltransferase porcupine suppresses cell growth and downregulates the Wnt/beta-catenin pathway in gastric cancer. Oncol Lett 5, 1719–1723.

Moorman, A., Webb, S., Brown, N.A., Lamers, W., and Anderson, R.H. (2003). Development of the heart: (1) formation of the cardiac chambers and arterial trunks. Heart 89, 806–814.

Muller, F.J., Brandl, B., and Loring, J.F. (2008). Assessment of human pluripotent stem cells with PluriTest. In StemBook (Cambridge (MA)).

Newman, A.M., Liu, C.L., Green, M.R., Gentles, A.J., Feng, W., Xu, Y., Hoang, C.D., Diehn, M., and Alizadeh, A.A. (2015). Robust enumeration of cell subsets from tissue expression profiles. Nature methods 12, 453–457.

Ng, W.A., Doetschman, T., Robbins, J., and Lessard, J.L. (1997). Muscle isoactin expression during in vitro differentiation of murine embryonic stem cells. Pediatric research 41, 285–292.

Panopoulos, A.D., D’Antonio, M., Benaglio, P., Williams, R., Hashem, S.I., Schuldt, B.M., DeBoever, C., Arias, A.D., Garcia, M., Nelson, B.C., et al. (2017). iPSCORE: A Resource of 222 iPSC Lines Enabling Functional Characterization of Genetic Variation across a Variety of Cell Types. Stem Cell Reports 8, 1086–1100.

Patel, S., Bonora, G., Sahakyan, A., Kim, R., Chronis, C., Langerman, J., Fitz-Gibbon, S., Rubbi, L., Skelton, R.J.P., Ardehali, R., et al. (2017). Human Embryonic Stem Cells Do Not Change Their X Inactivation Status during Differentiation. Cell Rep 18, 54–67.

Pau, G., Fuchs, F., Sklyar, O., Boutros, M., and Huber, W. (2010). EBImage--an R package for image processing with applications to cellular phenotypes. Bioinformatics 26, 979–981.

Pedregosa, F., Varoquaux, G., Gramfort, A., Michel, V., Thirion, B., Grisel, O., Blondel, M., Prettenhofer, P., Weiss, R., Dubourg, V., et al. (2011). Scikit-learn: Machine Learning in Python. J Mach Learn Res 12, 2825–2830.

Perez-Pomares, J.M., de la Pompa, J.L., Franco, D., Henderson, D., Ho, S.Y., Houyel, L., Kelly, R.G., Sedmera, D., Sheppard, M., Sperling, S., et al. (2016). Congenital coronary artery anomalies: a bridge from embryology to anatomy and pathophysiology--a position statement of the development, anatomy, and pathology ESC Working Group. Cardiovasc Res 109, 204–216.

Robinson, M.D., McCarthy, D.J., and Smyth, G.K. (2010). edgeR: a Bioconductor package for differential expression analysis of digital gene expression data. Bioinformatics 26, 139–140.

Robinson, M.D., and Smyth, G.K. (2008). Small-sample estimation of negative binomial dispersion, with applications to SAGE data. Biostatistics 9, 321–332.

Sahakyan, A., Plath, K., and Rougeulle, C. (2017). Regulation of X-chromosome dosage compensation in human: mechanisms and model systems. Philos Trans R Soc Lond B Biol Sci 372.

Scholzen, T., and Gerdes, J. (2000). The Ki-67 protein: from the known and the unknown. J Cell Physiol 182, 311–322.

Schwartzentruber, J., Foskolou, S., Kilpinen, H., Rodrigues, J., Alasoo, K., Knights, A.J., Patel, M., Goncalves, A., Ferreira, R., Benn, C.L., et al. (2018). Molecular and functional variation in iPSC-derived sensory neurons. Nature genetics 50, 54–61.

Stegle, O., Parts, L., Piipari, M., Winn, J., and Durbin, R. (2012). Using probabilistic estimation of expression residuals (PEER) to obtain increased power and interpretability of gene expression analyses. Nature protocols 7, 500–507.

Subramanian, A., Tamayo, P., Mootha, V.K., Mukherjee, S., Ebert, B.L., Gillette, M.A., Paulovich, A., Pomeroy, S.L., Golub, T.R., Lander, E.S., et al. (2005). Gene set enrichment analysis: a knowledge-based approach for interpreting genome-wide expression profiles. Proceedings of the National Academy of Sciences of the United States of America 102, 15545–15550.

Tan, A., Abecasis, G.R., and Kang, H.M. (2015). Unified representation of genetic variants. Bioinformatics 31, 2202–2204.

Tatler, A.L., Habgood, A., Porte, J., John, A.E., Stavrou, A., Hodge, E., Kerama-Likoko, C., Violette, S.M., Weinreb, P.H., Knox, A.J., et al. (2016). Reduced Ets Domain-containing Protein Elk1 Promotes Pulmonary Fibrosis via Increased Integrin alphavbeta6 Expression. J Biol Chem 291, 9540–9553.

Tohyama, S., Hattori, F., Sano, M., Hishiki, T., Nagahata, Y., Matsuura, T., Hashimoto, H., Suzuki, T., Yamashita, H., Satoh, Y., et al. (2013). Distinct metabolic flow enables large-scale purification of mouse and human pluripotent stem cell-derived cardiomyocytes. Cell stem cell 12, 127–137.

Tomoda, K., Takahashi, K., Leung, K., Okada, A., Narita, M., Yamada, N.A., Eilertson, K.E., Tsang, P., Baba, S., White, M.P., et al. (2012). Derivation conditions impact X-inactivation status in female human induced pluripotent stem cells. Cell stem cell 11, 91–99.

Wang, X., Moon, J., Dodge, M.E., Pan, X., Zhang, L., Hanson, J.M., Tuladhar, R., Ma, Z., Shi, H., Williams, N.S., et al. (2013). The development of highly potent inhibitors for porcupine. J Med Chem 56, 2700–2704.

Werner, J.C., and Sicard, R.E. (1987). Lactate metabolism of isolated, perfused fetal, and newborn pig hearts. Pediatric research 22, 552–556.

Witty, A.D., Mihic, A., Tam, R.Y., Fisher, S.A., Mikryukov, A., Shoichet, M.S., Li, R.K., Kattman, S.J., and Keller, G. (2014). Generation of the epicardial lineage from human pluripotent stem cells. Nature biotechnology 32, 1026–1035.

Zhao, M.T., Chen, H., Liu, Q., Shao, N.Y., Sayed, N., Wo, H.T., Zhang, J.Z., Ong, S.G., Liu, C., Kim, Y., et al. (2017). Molecular and functional resemblance of differentiated cells derived from isogenic human iPSCs and SCNT-derived ESCs. Proceedings of the National Academy of Sciences of the United States of America 114, E11111–E11120.

