## Supplemental Information for "Human iPSC gene signatures and X chromosome dosage impact response to WNT inhibition and cardiac differentiation fate"

### SUPPLEMENTAL TEXT

#### GSEA implicates ELK1 targets and genes on the X chromosome

To understand whether the transcriptomic differences between CM-fated iPSCs and EPDC-fated iPSCs were associated with alterations in specific pathways or cellular function, we performed a gene set enrichment analysis (GSEA) on 9,808 MSigDB gene sets [1, 2] using the 15,228 expressed autosomal genes in the 184 iPSCs (Table S16). We identified 22 gene sets that were significantly associated with iPSC cell fate, including enrichment in the 125 CM-fated iPSCs for transcription factor activity, DNA binding and ELK1 targets, and enrichment for the 59 EPDC-fated iPSCs for extracellular matrix (Figure 4A). These results show that EPDC-fated iPSCs were more likely predisposed to develop an extracellular matrix scaffold that is typical of fibroblasts and, in general, mesenchymal tissues [3]. To capture gene sets associated with expression differences on the X chromosome, we performed differential expression and GSEA on 113 female iPSC lines (87 CM-fated; 26 EPDC-fated). We identified 2 significant gene sets comprised of loci on the chromosome X (chrXp11 and chrXp22, Figure 4A, Table S16). Interestingly, chrXp11 includes the *ELK1* locus, suggesting that the overexpression of ELK1 targets may be due to an overall higher transcriptional activity of the *ELK1* locus. *PORCN*, whose function in WNT signaling activation is inhibited by IWP-2 at D3 through D5 differentiation [4-6], is located 800 kb downstream of ELK1, also in the chrXp11 locus. The chrXp22 locus includes the majority of genes (52/99, 52.5%) that are known to escape chromosome X inactivation [7, 8], and thus may potentially have varying X-linked gene dosage across female iPSCs. Overall, GSEA shows that genes differentially expressed between CM-fated and EPDC-fate iPSCs are involved in a variety of pathways including ELK targets and potentially associated with the X chromosome activation status.

#### Sex associated with iPSC differentiation outcome

To identify other iPSC factors potentially associated with differentiation outcome, we examined three characteristics of the 181 subjects in our study (sex, ethnicity and age) and passage of the iPSCs at D0 (Figure 4B, S14; Tables S1, 2). Analyzing the 125 CM-fated and 59 EPDC-fated iPSC lines with a general linear model, we found no association between differentiation outcome and age or ethnicity ($p > 0.8$; GLM, Z-test, Figure S12A,B; Table S17), but observed a significant association with sex ($p = 2.57 \times 10^{-5}$, GLM, Z-test; Figure 4B) and a trend for iPSC passage at D0 ($p = 0.069$, GLM, Z-test; Figure S12C). These data suggest that iPSCs derived from female subjects and iPSCs with higher passages at D0 had an increased predisposition for the CM fate. Furthermore, considering only the 193 completed differentiations (D25 iPSC-CVPC samples), we found that iPSC-CVPC samples derived from female subjects compared to those derived from males had significantly higher %cTnT values (mean = 83.0% and 77.7%, respectively for females and males, $p = 6.0 \times 10^{-4}$, Mann-Whitney U test; Figure S12D) and a higher fraction of CMs ($p = 6.46 \times 10^{-4}$, Mann-Whitney U test; Figure S12E). These results indicate that iPSCs derived from female subjects and, to a lesser extent, iPSCs that have spent more time in cell culture, have a greater inherent predisposition to differentiate towards the CM lineage.

### Female iPSCs with X chromosome reactivation associated with an CM fate

Given the observation that iPSCs from females have a greater potential to give rise to CMs, we asked if X chromosome inactivation (Xi) status was associated with CM or EPDC fate. Using RNA-seq data generated from the 113 female iPSCs, we evaluated allele specific effects (ASE) of X chromosome and autosomal genes (Table S18). We defined the strength of ASE for each gene as the fraction of RNA transcripts that were estimated to originate from the allele with higher expression (hereto referred to as allelic imbalance fraction, AIF). We observed that, while AIF in autosomal genes was close to 0.5, indicating both alleles were equally expressed (Figure 4C), AIF on the X chromosome in iPSCs tended to be bimodal, with a larger fraction of genes showing monoallelic expression (AIF ~1.0). We observed that AIF was significantly less in the 87 CM-fated female iPSCs compared with the 26 EPDC-fated female iPSCs ($p = 0.011$, Mann-Whitney U test, Figure 4D) and that this difference in AIF became even more pronounced in the derived CM and EPDC samples ($p = 4.81x10^{-6}$, Mann-Whitney U test) (Figure 4E). These observations are consistent with previous studies showing that X chromosome reactivation does not reverse during differentiation [9, 10]. These observations show that female iPSCs with an eroded X chromosome were associated with a CM fate.

Since we observed differences in X chromosome reactivation state between CM-fated and EPDC-fated female iPSCs, we next asked if the two GSEA X chromosome associated intervals (chrXp22 and chrXp11; Figure 4A) showed corresponding allelic imbalance trends. We plotted AIF differences, where a positive AIF difference indicates X chromosome reactivaton in the 26 EPDC-fated iPSCs and a negative in the 87 CM fated-iPSCs (Figure S13A). We observed that distinct regions across the X chromosome were differentially eroded in the EPDC-fated versus CM-fated iPSCs. In particular, chrXp22 showed X reactivation in CM-fated iPSCs ($p = 6.31 \times 10^{-3}$, Mann Whitney U), with both escape ($p = 0.020$, Mann Whitney U) and non-escape genes ($p = 0.023$, Mann Whitney U) showing evidence of reactivation (Figure S13B-D). As chrXp22 contains more than half of escape genes on the X chromosome, this observation confirms that increased X reactivation in CM-fated iPSCs results in increased expression of both escape and non-escape genes. As GSEA identified genes on chrXp11 to be overexpressed in CM-fated iPSCs, the lack of X reactivation in this interval ($p = 0.28$, Mann Whitney U) suggests alternative regulatory mechanisms may also alter gene expression levels on the X chromosome. Overall, these results suggest that differential X chromosome reactivation as well as other altered regulatory effects in iPSCs may contribute to cardiac lineage fate.

### SUPPLEMENTARY FIGURES

#### Figure S1: Optimization of iPSC-CVPC differentiation protocol

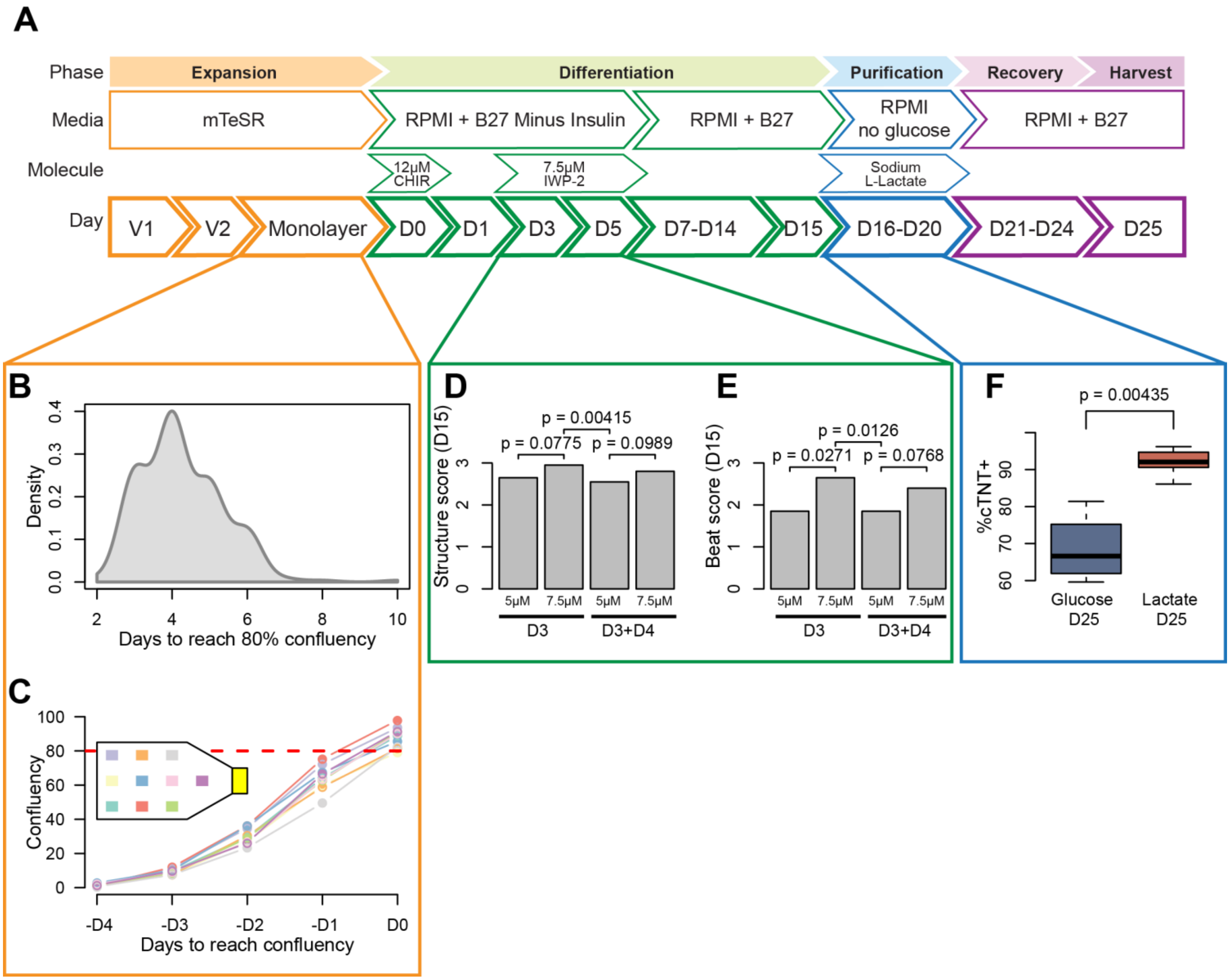

(A) Schematic of differentiation protocol. To achieve a large-scale derivation of iPSC-derived cardiovascular progenitor cells (iPSC-CVPCs), we optimized existing small molecule protocols to increase throughput and efficiency. In order to use existing protocols for iPSC-CVPC differentiation in a large-scale experiment, sources of experimental variability between iPSC lines need to be minimized. We optimized several steps of protocol, including automating detection iPSC monolayer confluency (orange box), optimizing the IWP-2 concentration (green box), and incorporating lactate selection (blue box), were these together allowed the performance of a large-scale differentiation of iPSC-CVPCs, minimizing the experimental differences between lines.

(B) Density plot showing distribution of days recorded from 253 iPSC samples to reach 80% confluency. It was observed that 75-85% iPSC monolayer confluence at day 0 (D0), which marks the initiation of differentiation by WNT activation, yields the most efficient differentiations [4, 6]; however, iPSCs have variable growth rates, and therefore it would be difficult to consistently achieve this confluency across hundreds of lines.

(C) Confluency levels from one line (2_3) measured from ten sections of T150 flask. Due to observed variability in iPSC growth rates, we developed ccEstimate, an automated tool that determines confluency by processing images from multiple locations in three T150 flasks over a period of at least 72 hours, and then estimates when a particular iPSC line will reach an average confluency of 80% based on its growth rate (Figure S2). Circles represent measured values. The points at D0 were obtained based on the ccEstimate algorithm's predictions.

(D, E) Effects of IWP-2 concentration (5.0µM or 7.5µM) given on D3 or D3 and D4 on (D) structure score and (E) beat score (Table S20). We optimized WNT inhibition at D3, which is required for robust iPSC-CVPC differentiations by testing two concentrations of IWP-2 (5µM and 7.5µM) both with and without a media change between D3 and D4. We differentiated one iPSC line (iPSCORE_2_3_iPSC_C5_P13) under each of the four IWP-2 conditions, and observed that 7.5 µM IWP-2 without a media change between D3 and D4 resulted in iPSC-CVPCs with the thickest structures and strongest beating (structure score: $p = 0.00415$, beat score: $p = 0.0126$; Paired t test) (Table S20). P-values were calculated using paired t test.

(F) Effects of metabolic purification of iPSC-CVPCs by lactate and glucose. To examine the efficacy of using lactate for iPSC-CVPC metabolic purification [4, 11, 12], we tested lactate and glucose at D16 in three different iPSC-CVPC lines (2_3, 8_2, and 3_2), and found that lactate resulted in significantly purer iPSC-CVPC populations (93.95% vs. 68.55%; $p = 0.00435$). P-values were calculated using Mann-Whitney U test.

**Figure S2: Examples of cell confluency detection using the ccEstimate algorithm**

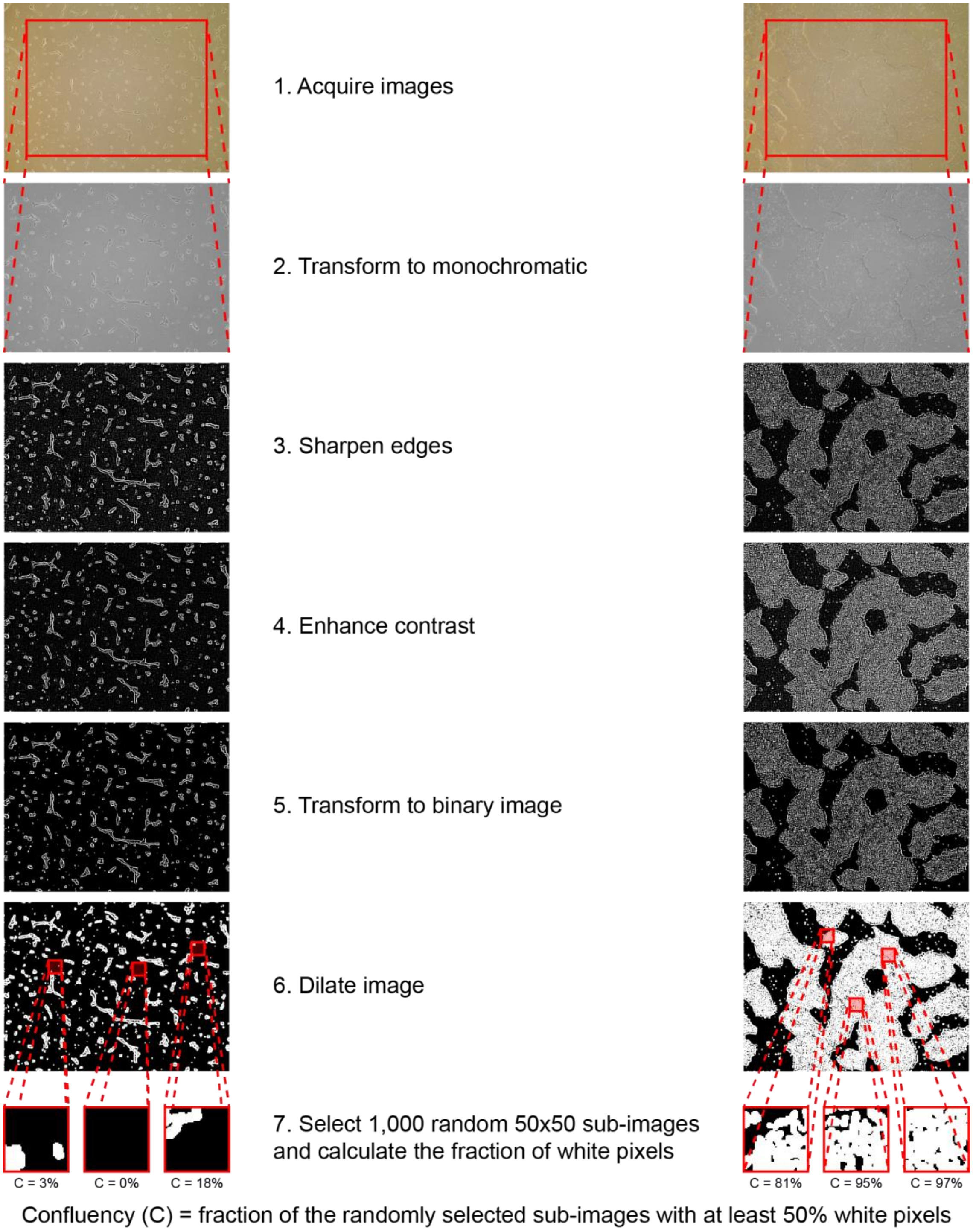

The image shows two examples of the ccEstimate algorithm that we developed to automatically determine the confluency, which includes 7 steps: 1) image acquisition, 2) monochromatic transformation, 3) edge sharpening, 4) contrast enhancement, 5) binary image transformation, 6) image dilation, and 7) fraction of white pixel calculations from 1,000 sub-images. On the left column of images, a low confluency iPSC sample is shown (UDID016 at D1), while on the right column of images, a high confluency sample (UDID016 at D4) is shown.

**Figure S3: Immunofluorescence staining of D30 iPSC-CVPCs**

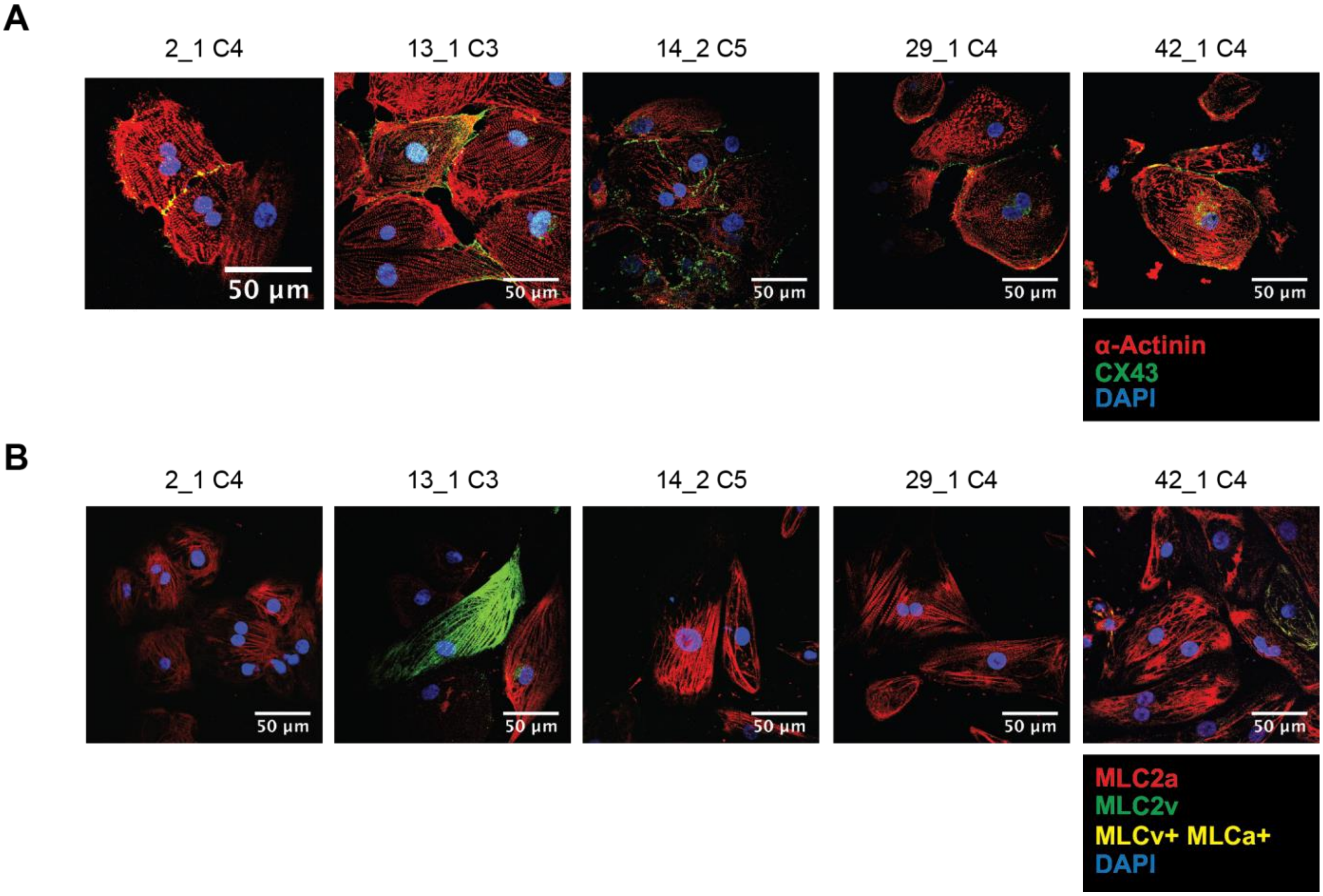


(A) Immunofluorescence staining of five iPSC-CVPC lines with IF markers DAPI (blue), ACTN1 (red), and CX43 (green). (B) Immunofluorescence staining of five iPSC-CVPC lines with IF markers DAPI (blue), MLCa+ (red), and MLCv+ (green), and MLCv+ MLCa+ (yellow).

**Figure S4: Filtering doublets from and selection of k-means clustering k in scRNA-seq**

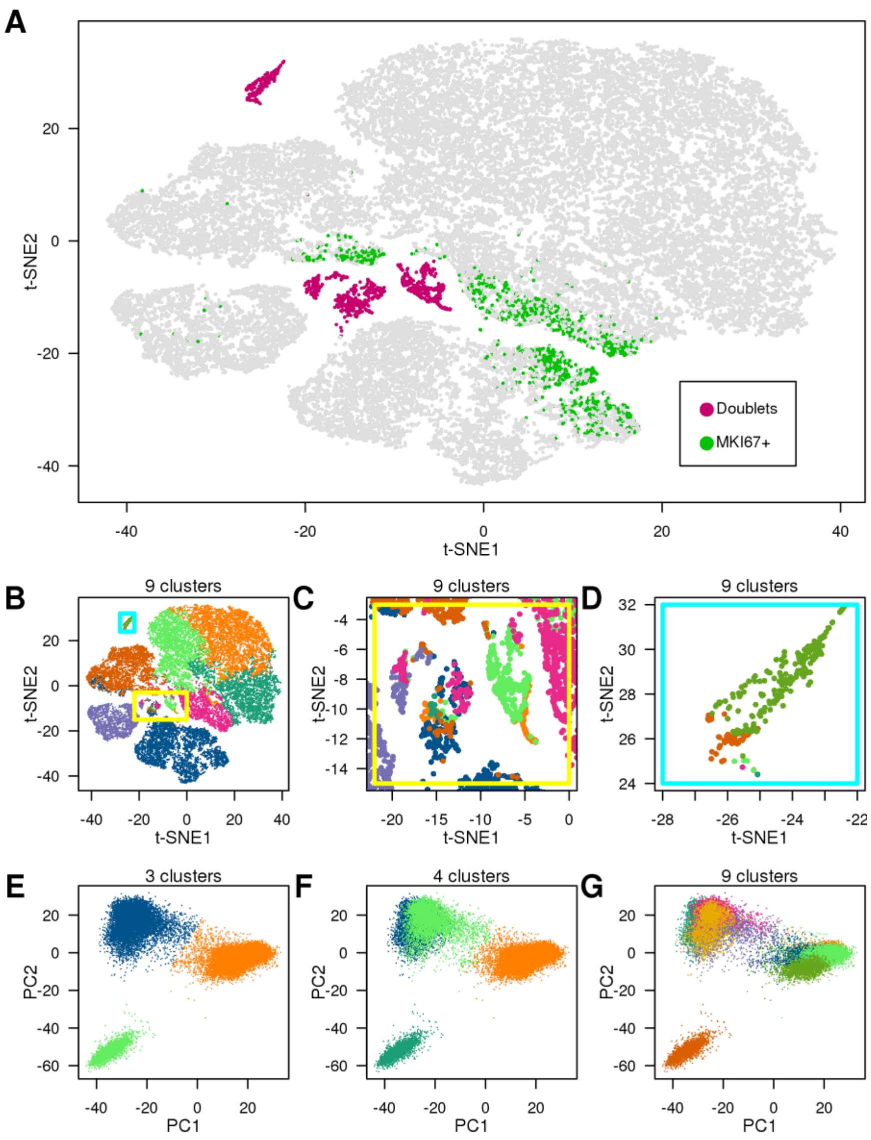


(A, B) t-SNE plot of gene expression from 36,839 cells from 8 iPSC-CVPC and 1 ESC. We removed 1,934 cells from the scRNA-seq analysis (Figure 2), including cells that were visually identified as being doublets (Figure S4 B-D) (pink) and actively dividing cells with (>2 UMI MK167, green). (B) Cells are colored by k-means clusters with k = 9 cluster assignment. Cells that clustered together in the t-SNE plot, but were assigned to multiple different clusters were considered as doublets. Doublets are highlighted in the yellow and cyan box. (C) Zoomed in region of the yellow box (Figure S4B). (D) Zoomed in region of the cyan box (Figure S4B). (E-G) PCA of gene expression from 36,839 cells from 8 iPSC-CVPC and 1 ESC colored by k-means clustering: (E) k = 3, (F) k=4; and (G) k = 9. The PCA shows that three cell populations are present and thus we used three clusters (k = 3) for all scRNA-seq analysis. In summary, we analyzed 34,905 single cells assigned to three cell populations.

**Figure S5: Distribution of single cells across the three cell populations for the nine analyzed samples**

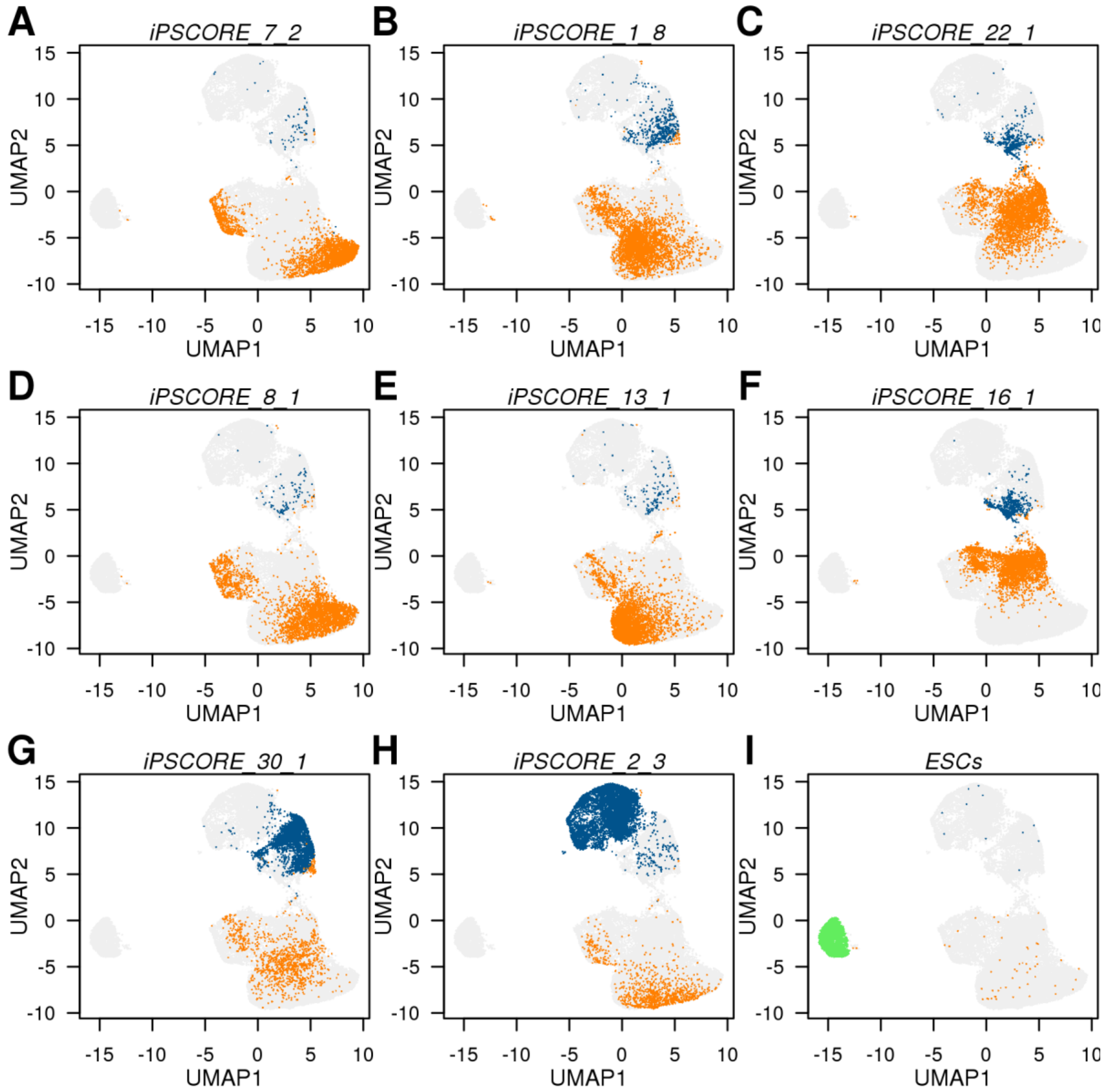


(A-I) scRNA-seq UMAP plots from 34,905 cells showing the distribution of the nine analyzed samples (8 iPSC-CVPCs lines and one ESC line) across the three different clusters. Each plot only shows cells associated with one sample. Population 1 = orange; Population 2 = blue; Population 3 = green.

**Figure S6: Expression levels for marker genes**

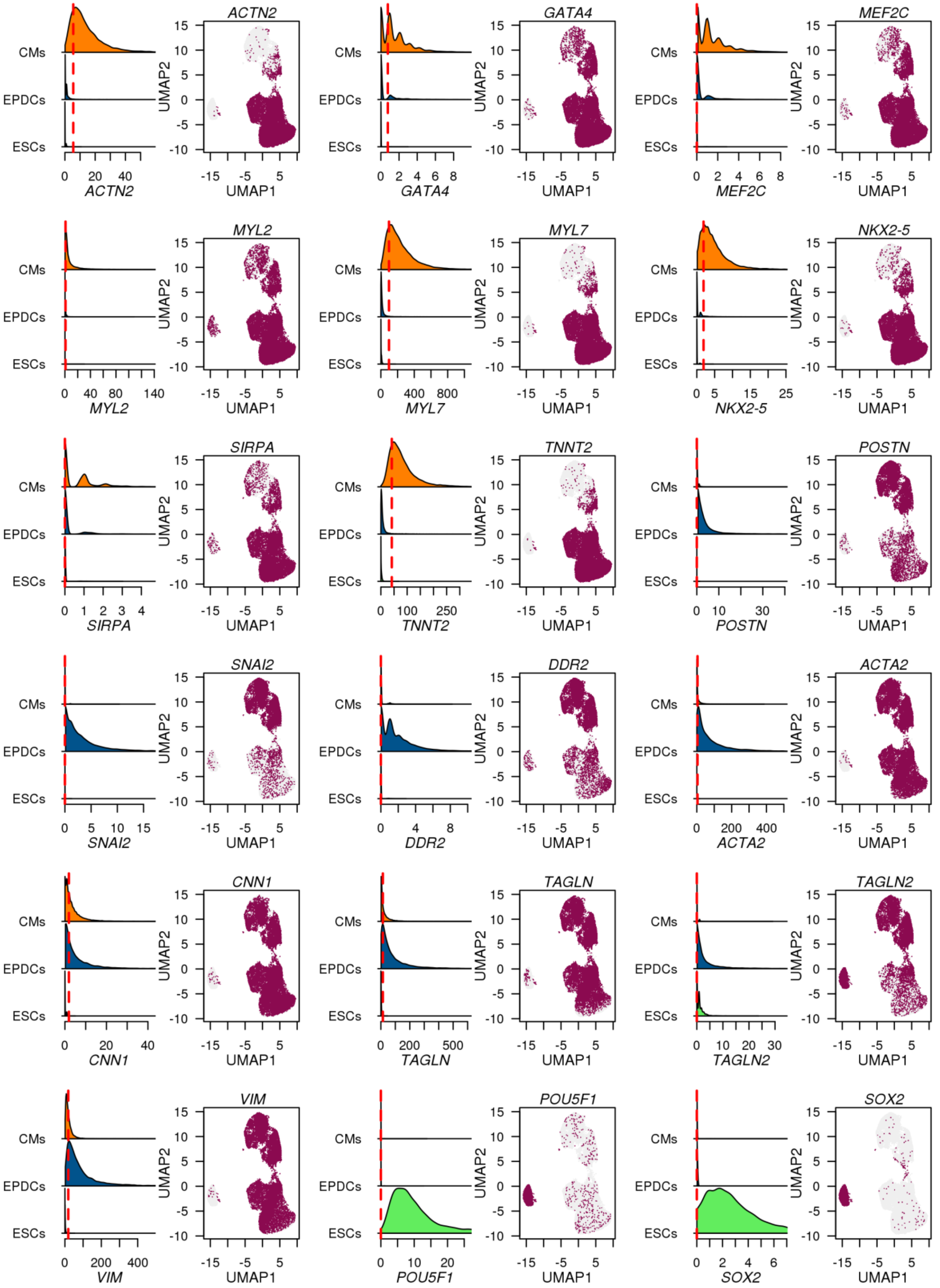

For each gene in Figure 2D, density plots show the gene expression distribution across all cells associated with each cell population (Population 1 = orange; Population 2 = blue; Population 3 = green.). Red dashed line represents the median. UMAP  plots from 34,905 cells show in maroon all the cell with expression higher than the median expression across the three populations.

**Figure S7: Volcano plots showing differential expression at different CM population thresholds**

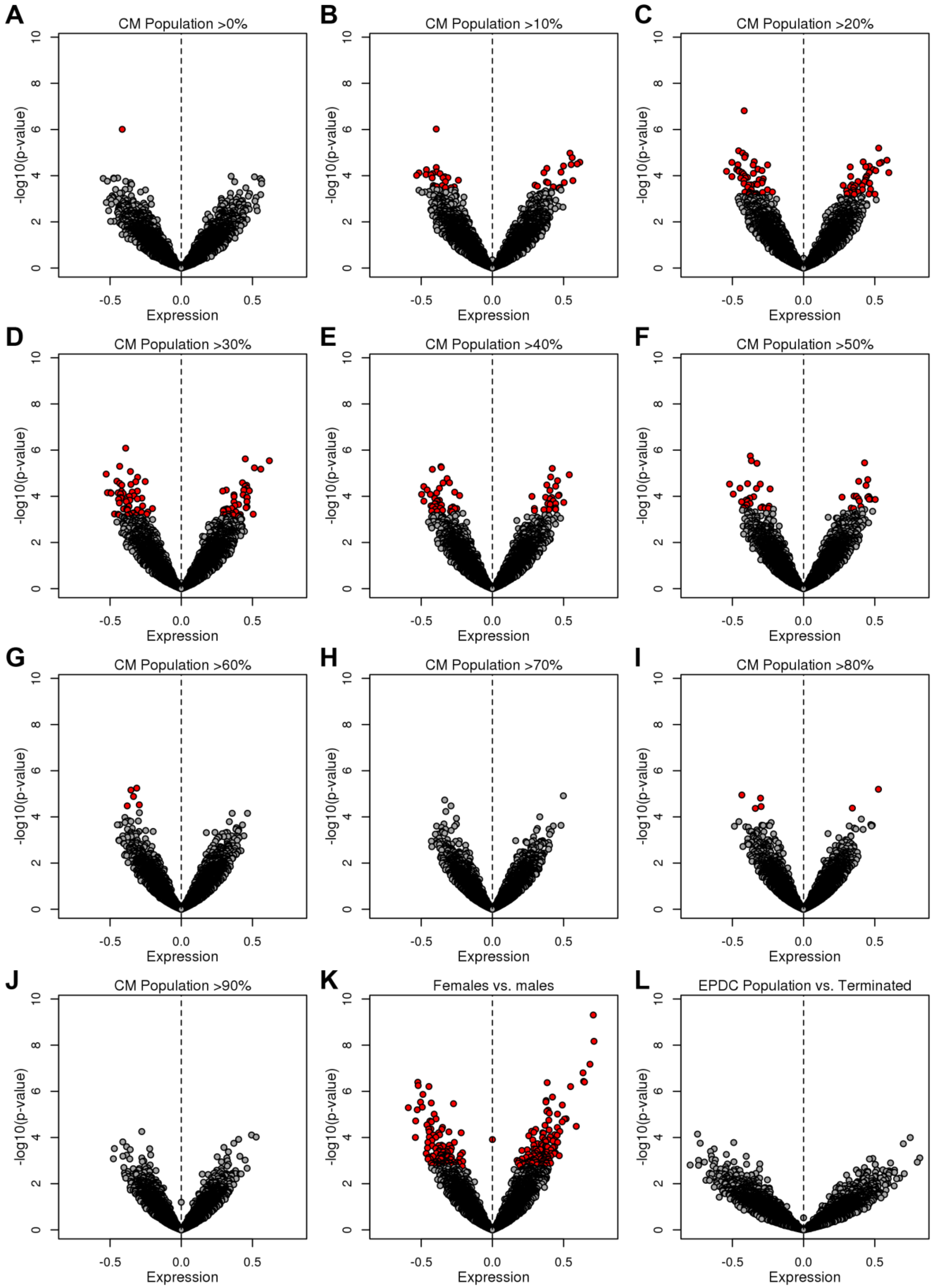

(A-J) Volcano plots displaying data used to determine fraction of Population 1 in iPSC-CVPC samples (i.e. >30% CMs) to define iPSCs as having either a CM fate or an EPDC fate. Here, we show the number of differentially expressed genes between the two groups of iPSCs (CM or EPDC fate) at varying thresholds of Population 1: >0%, 10%, 20%, 30%, 40%, 50%, 60%, 70%, 80%, 90%. The data for genes differentially expressed at each of these thresholds is shown in Table S9. At the 30% threshold, there were the greatest number of significantly differentially expressed genes. EPDC fated iPSCs consisted of those with completed iPSC-CVPC differentiations (i.e. reached D25) with >70% Population 2 and iPSCs with differentiations that were terminated before D25. (K) We examined genes that were differentially expressed in the iPSCs between female and male samples, and removed genes that overlapped with the set of differentially expressed genes in (D). (L) We did not observe significant expression differences between iPSCs that differentiated to EPDCs (>70% Population 2) and iPSCs whose differentiations were terminated before D25. (A-L) Mean difference in expression (X axis) and p-value (Y axis, t-test) of each threshold are shown. Significant genes are indicated in red and thresholds are indicated above each plot. Based on these analyses, we defined iPSC-CVPCs with ≥30% Population 1 as CMs and those with <30% Population 1 as EPDCs and the matched iPSCs respectively as CM-fated and EPDC-fated.

**Figure S8: Comparison between the observed number of signature genes and random expectation**

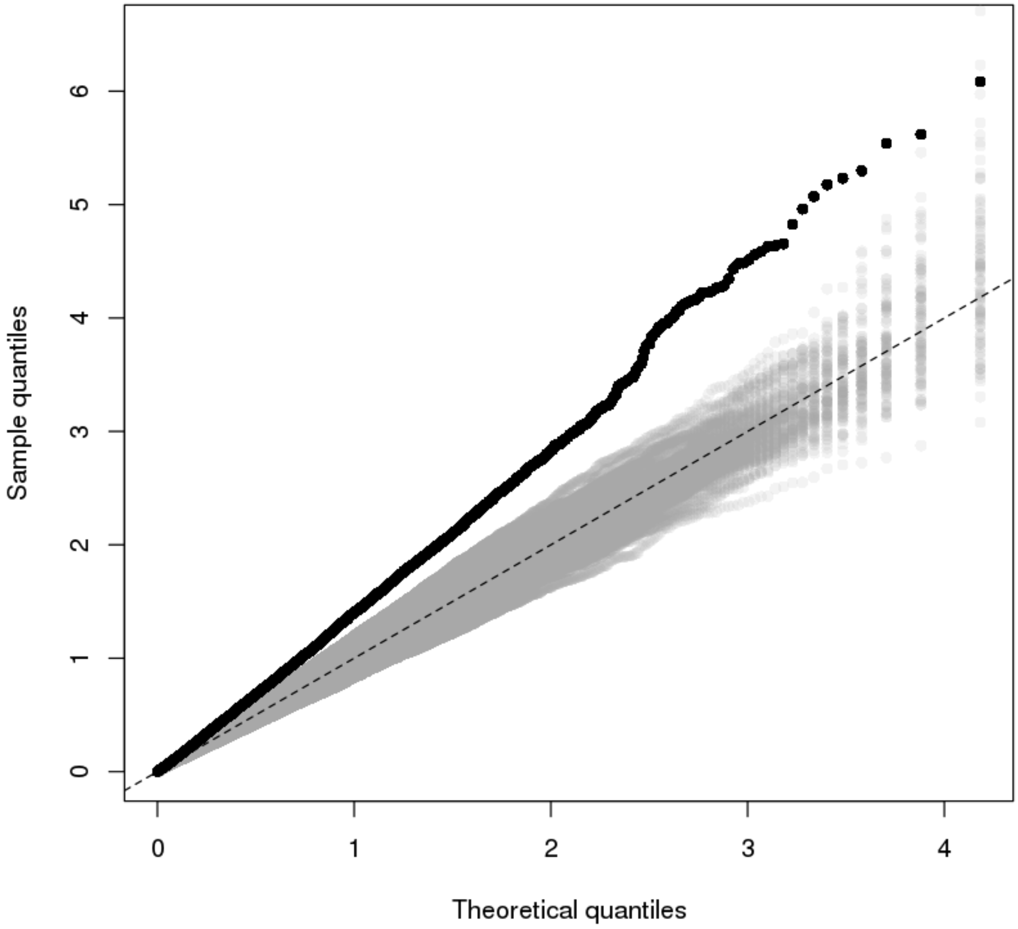


A QQ plot showing that the observed p-value distribution (black) was substantially different than random expectation. To determine if the identification of 84 signature genes that were significantly differentially expressed between CM-fated and EPDC-fated iPSCs was higher than random expectation, we shuffled the assignments of the 184 iPSC RNA-seq samples to differentiation fates (125 CM and 59 EPDC) 100 times. For each shuffle, we performed differential expression analysis and obtained the number of genes that were significantly differentially expressed (gray).

**Figure S9: Expression levels of 91 signature genes**

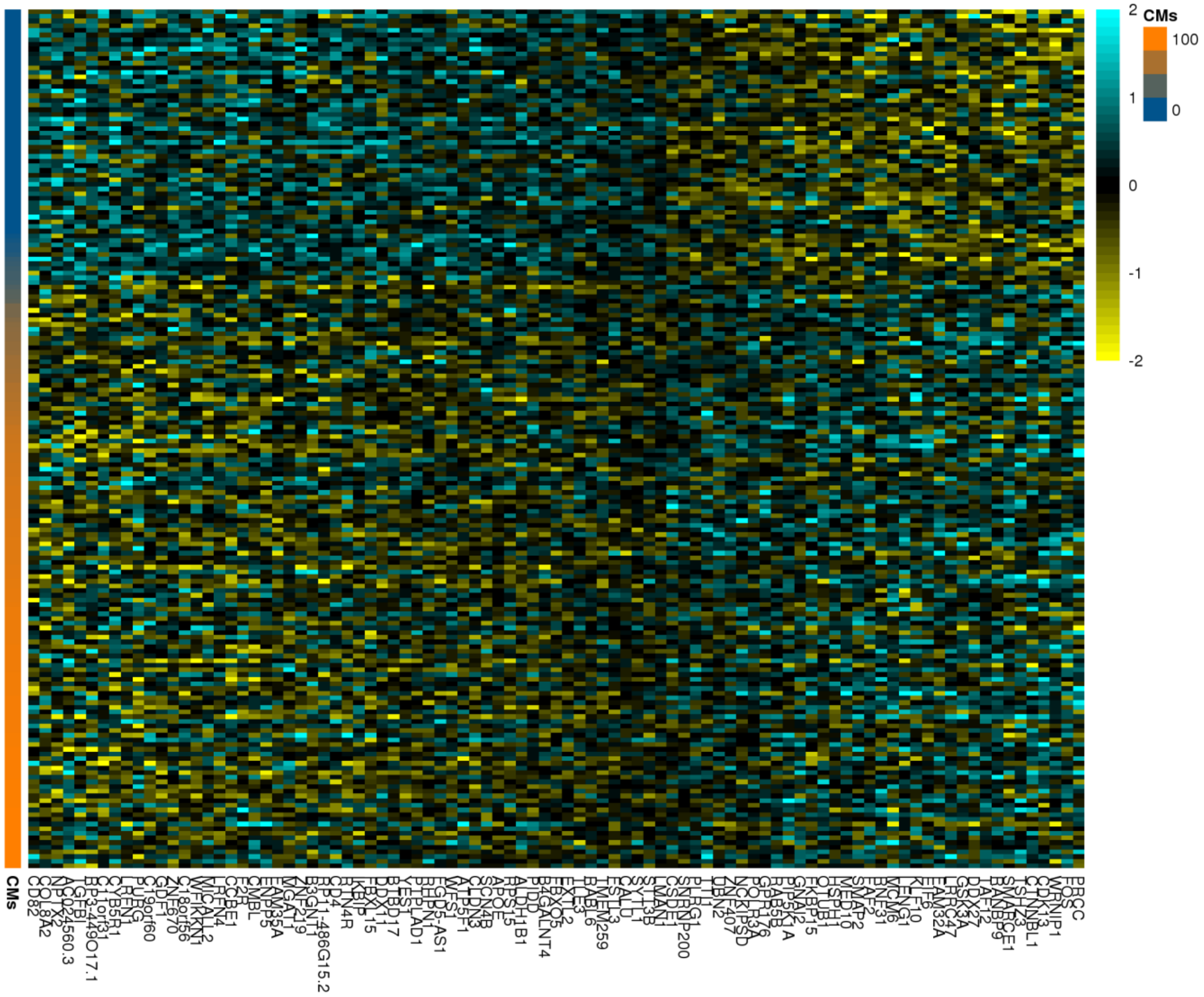


Heatmap showing the expression levels of the 91 signature genes differentially expressed between iPSCs with CM fate and iPSCs with EPDC fate. Each row represents an iPSC sample. The “CMs” scale represents the %CM population (Population 1) in the associated iPSC-CVPC samples for each iPSC.

**Figure S10: Correlation between the 91 signature genes differentially expressed between iPSCs with a CM fate and iPSCs with an EPDC fate**

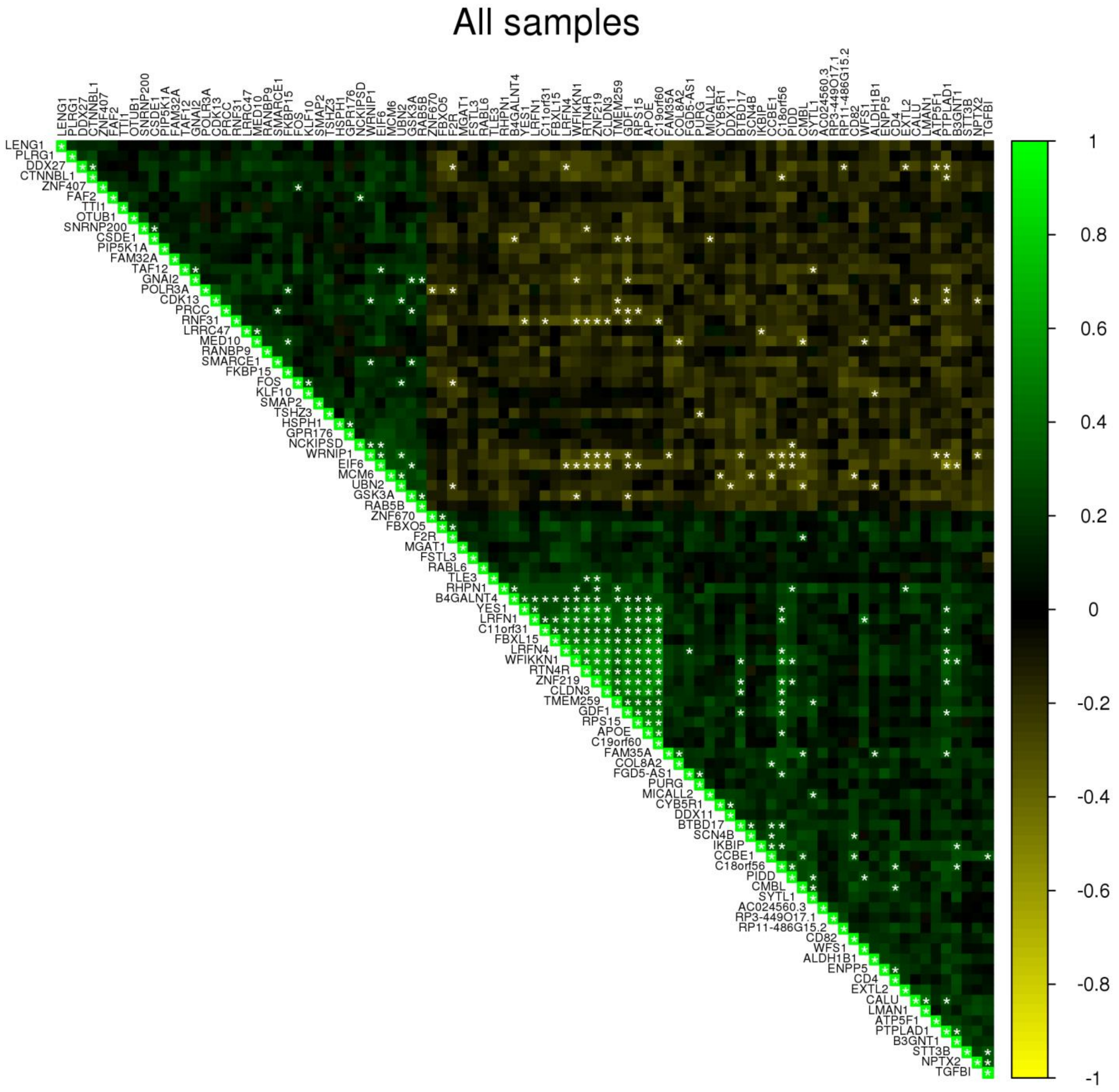


Heatmap showing the correlation of expression levels in the iPSCs with a CM fate versus the iPSCs with an EPDC fate for the 91 signature genes. White stars show significant correlations (Bonferroni p-value < 0.05).

**Figure S11: Associations between genetic variation and differentiation outcome**

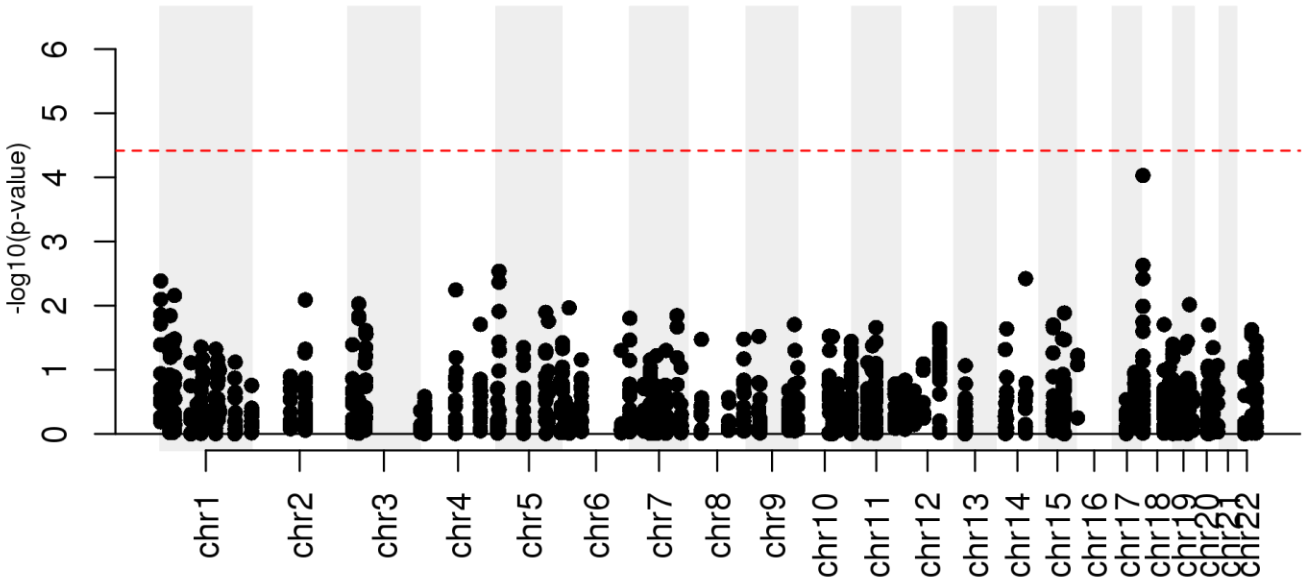


(G) Manhattan plot showing the association between genetic variation for SNPs that are eQTLs for the 91 signature genes and differentiation outcome (measured as % CM population in iPSC-CVPCs). Red dashed line shows p-value = 0.05 adjusted using Bonferroni's method.

**Figure S12: X chromosome inactivation in iPSCs**

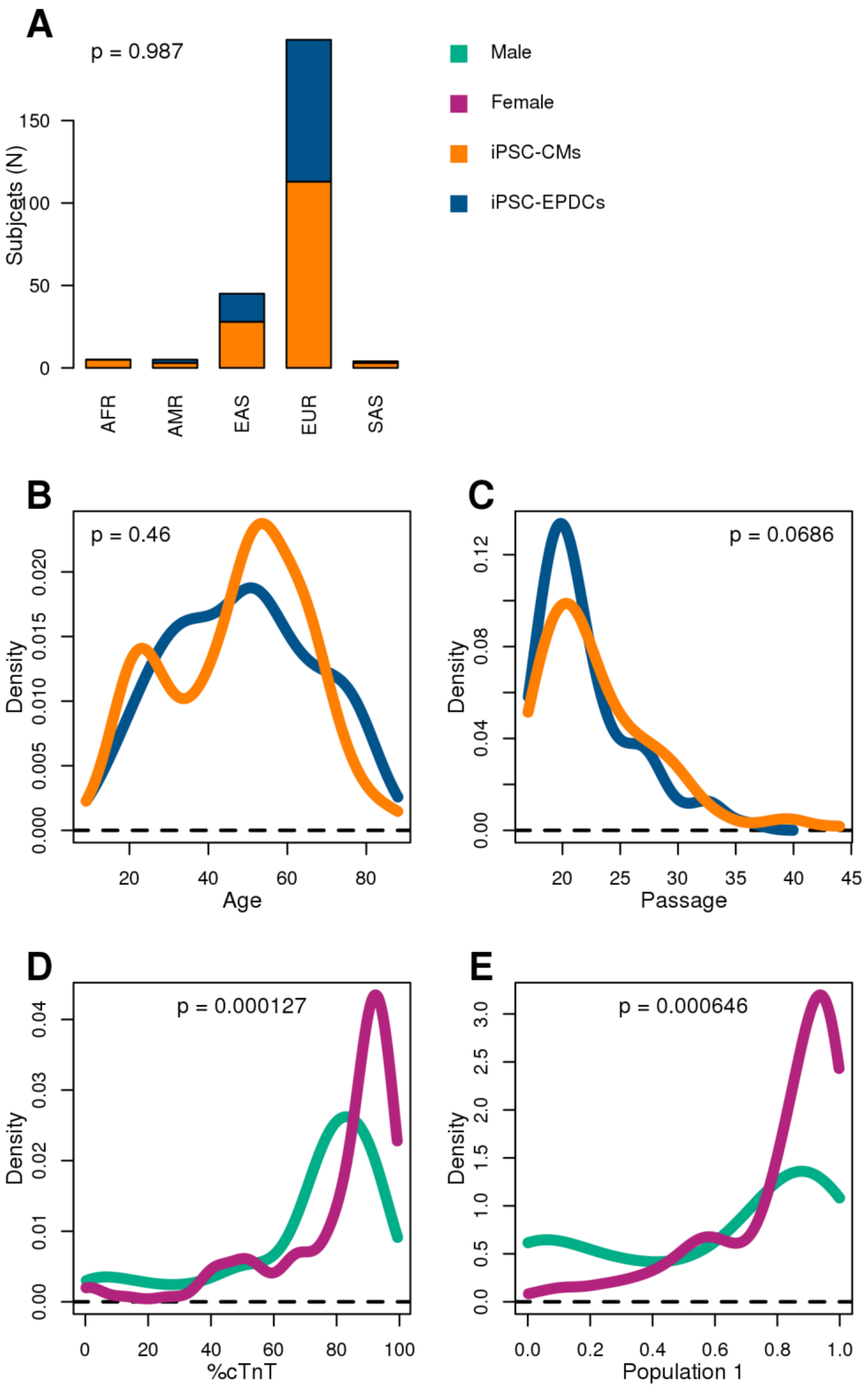


(A-C) Associations between differentiation outcome (orange: iPSC-CVPC samples with CM fraction > 30%; blue: with EPDC fraction > 70%) and (A) ethnicity (most similar superpopulation from the 1000 Genomes Project), (B) age at enrollment, and (C) passage at monolayer (D0). (A) is shown as barplots; (B,C) are shown as density plots. P-values were calculated using Z-test (glm function in R).

(D-E) Density plots showing the association between sex (teal: males; magenta: females) and (D) %cTnT, and (E) fraction of CM population for 191 iPSC lines. P-values were calculated using Mann-Whitney U test.

**Figure S13: Allelic imbalance fraction from inactive and escape genes in Xp22**

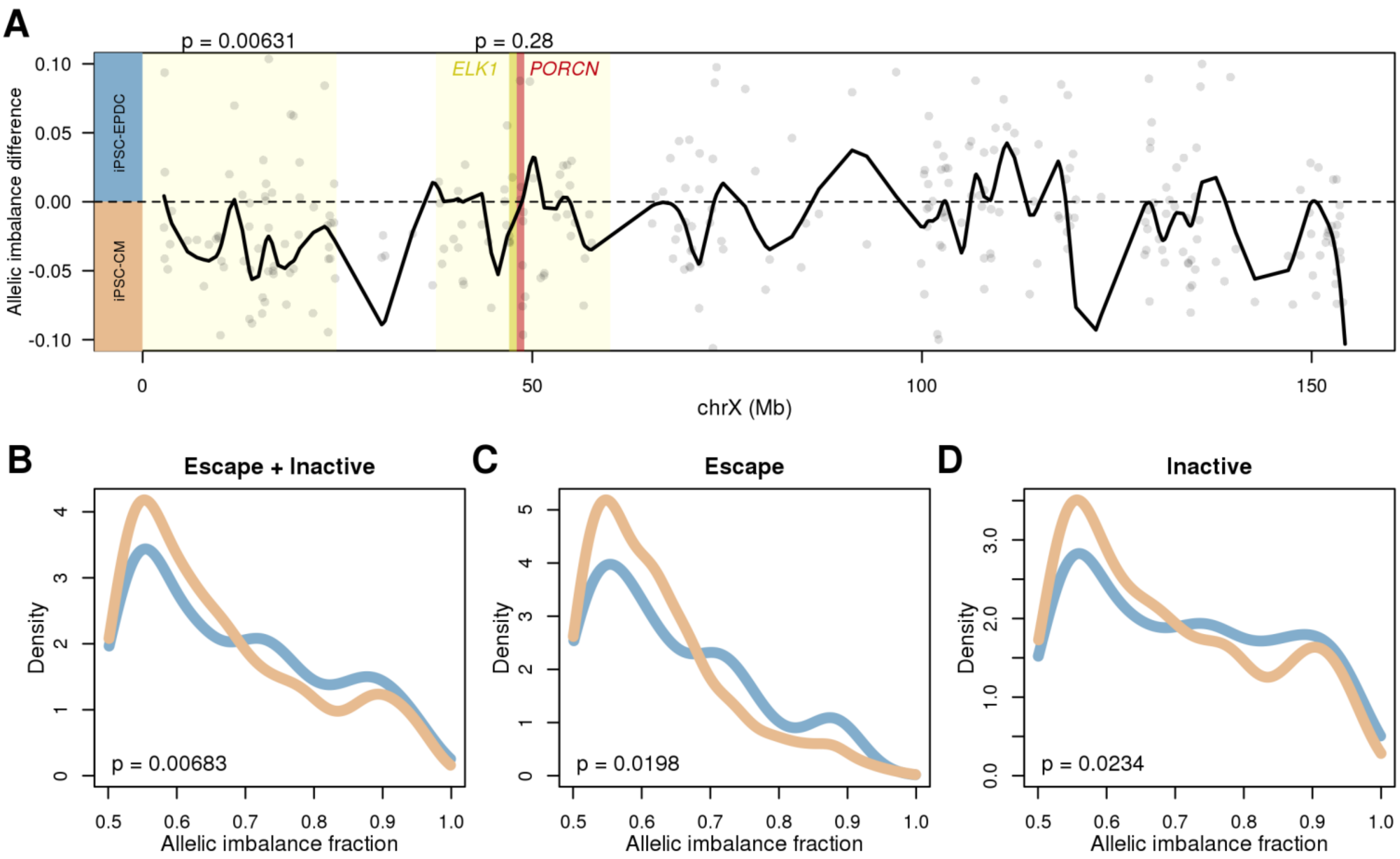


(A) Allelic imbalance difference between CM-fated and EPDC-fated iPSCs. The dots represent each gene on chrX, while the black solid line corresponds to the smoothed interpolation of differences for all the genes. Xp22 and Xp11 loci on the chrX G-banding ideogram are highlighted in blue, as well as ELK1 (yellow) and PORCN (red). P-values above each locus indicate the difference in allelic imbalance between CM fated and EPDC fated iPSCs in each locus (Mann Whitney U test).

(B-D) Density plots showing the allelic imbalance differences in chrX genes on the Xp22 loci in female samples between iPSC lines with CM fate (light blue) and EPDC fate (light orange) differentiations. Allelic balances compared in Xp22 are from all genes in the region (B), escape genes (C), and inactive genes (D). P-values were calculated using Mann Whitney U test.

### TABLE LEGENDS

**Table S1. Subject information for participants in iPSCORE for which iPSCs were used for iPSC-CVPC differentiation.**

iPSCORE_ID indicates family and individual number (e.g. iPSCORE_family#_individual#). Subject_UUID is an assigned Universal Unique Identifier (UUID) for the subject. Family_ID classifies the subject by family to identify related family members. Columns D-G represent the twin and parent information for each subject, as included in dbGap (phs001325.v1.p1; phs000924.v1.p1) as part of the iPSCORE Resource: Twin_ID_dbgap identifies the dbGap id if the subject is a twin; Twin_type_dbGap indicates the type of twin (MZ = monozygotic; DZ = dizygotic) if the subject is a twin; Father_subect_ID_dbGap indicates the subject_UUID of the father of the subject if part of the iPSCORE resource; Mother_subect_ID_dbGap indicates the subject_UUID of the mother of the subject if part of the iPSCORE resource. Sex and Age_at_enrollement of the subject are shown. Ethnicities (Self-reported race/ethnicity, Recorded_Ethnicity_Grouping, and Most_similar_1KGP_population) are recorded as described by Panopoulos et al. [13].

**Table S2. Table linking identifiers for iPSCORE participants with iPSC-CVPC differentiations and metrics of differentiation outcome.**

Unique Differentiation IDentifier (UDID) is a unique digit assigned for each attempted iPSC-CVPC differentiation. iPSCORE_ID indicates family and individual number (e.g. iPSCORE_family#_individual#). Subject_UUID is an assigned Universal Unique IDentifier (UUID) for the subject. iPSC_iPSCORE_ID is the iPSC line identifier submitted to dbGap (phs001325.v1.p1), which indicates clone and passage of iPSC. iPSCORE_resource indicates by TRUE or FALSE if this line is one of the 222 lines described by Panopoulos et al. [13]. iPSC_ID is the iPSC line identifier. iPSC_passage_at_monolayer (D0) is reported. D_to_D0 describe how many days the iPSC line was cultured to achieve 80% confluency before initiation of differentiation. If the UDID was harvested on D25 (Column I), the harvest density (Column J), number of cryovials frozen (Column K), and measured %cTNT+ by FACS (Column L) is reported. Successful_iPSC_CM_differentiation indicates if the iPSC-CVPC sample was harvested at D25 (e.g. not prematurely terminated). Cluster_1 indicates the estimated composition of population 1 (cardiomyocyte population) for each sample with RNA-seq (see Supplemental Table 3) and the estimated_cell_type (Column O) indicates samples with ≥30% population 1 as CM and samples with <30% population 1 as EPDC.

**Table S3. Table describing the number of lines and subjects for each attempted differentiation**

For each cell type: iPSC and derived iPSC-CVPCs (both terminated prior to D25 and D25), the number of differentiations performed (Column B) are given. For these differentiations, the number of unique lines used (Column C) from the number of unique subjects used (Column D) is provided.

**Table S4 Antibodies used for FACS and immunofluorescence**

This table describes the antibodies (Column A) and clone (Column B) used for FACS and immunofluorescence experiments. Catalog numbers (Column C), brand (Column D), dilution (Column E), time of staining in minutes (Column F), and temperature of staining (Column G) is indicated for each antibody.

**Table S5. Table linking identifiers for iPSC and iPSC-CVPC genomic data.**

UDID is given if the iPSC or iPSC-CVPC genomic data were collected during an attempted differentiation, indicated by UDID (Column A). Subject_UUID (Column B) is an assigned Universal Unique Identifier (UUID) for the subject. Cell (Column C) indicates the stage for which the genomic data was generated (iPSC or iPSC-CVPC). Genomic data UUIDs are given in Columns D-E, including rna_assay_uuid (bulk RNA-seq; Column D), scrna_assay_uuid (scRNA-seq; Column E). Estimated cellular composition from CIBERSORT of populations 1-3 is given in Columns F-H.

**Table S6. Generated molecular data**

For each cell type (iPSC and iPSC-CVPC) (Column A) and for each assay for which molecular data was generated (RNA-seq and scRNA-seq), the number of data samples (Column C), from the number of unique lines (Column D), and from the number of unique subjects (Column E) are given.

**Table S7: scRNA-seq features for each sequenced single cell**

For each of the 34,905 cells with scRNA-seq data, the table shows iPSCORE subject ID (Column A) and UUID (Column B), barcode (Column C), associated population (Column D), and coordinates on the t-SNE plot (Columns E, F). For the H9 ESC line sample, which is not included in iPSCORE, iPSCORE ID and Subject UUID are labeled as “ESCs”. This table is ordered on population (e.g. clusters 1, 2, 3).

**Table S8: Overexpressed genes in each scRNA-seq population**

For each of 34,528 genes (Columns A, B) with at least one transcript detected in the scRNA-seq samples, the mean UMI counts, log2 fold change, and FDR-adjusted p-value is shown for each population. The last column indicates the 150 genes used as input for Cibersort.

**Table S9: Differential expression between iPSCs differentiated to CMs and iPSCs differentiated to EPDCs using multiple thresholds**

We used 10 different thresholds to divide iPSCs based on their %CM population detected using Cibersort (Figure S8). For each threshold (Columns B-M) (>0%, 10%, 20%, 30%, 40%, 50%, 60%, 70%, 80%, 90%; males and females; EPDC fate vs. Terminated), differential expression between the samples passing and not passing the threshold was calculated using t-test. P-values of differentially expressed genes calculated by t-test are shown for all 15,228 human genes (Column A) expressed in iPSC-CVPCs. For further analyses, we used the 30% threshold, because it maximized the differences between CMs and EPDCs, i.e. the largest number of differentially expressed genes was obtained using 30% threshold.

**Table S10: Differential expression analysis using 30% CM population as threshold**

In this table, we report the differential expression analysis performed between iPSC samples that differentiated to CMs and iPSC samples that differentiated to EPDCs (15,228 autosomal genes, labeled as "All iPSC samples" in column H), as well as the differential expression analysis performed between female iPSC samples that differentiated to CMs and female iPSC samples that differentiated to EPDCs (15,398 expressed genes located on autosomes and on chromosome X, labeled as "Female samples" in column H). For each gene (Columns A, B), shown are: 1) the mean normalized expression across the iPSC samples that differentiated to CMs (column C), 2) the mean normalized expression across the iPSC samples that differentiated to EPDCs (column D), 3) the difference between mean normalized expressions (column E), 4) the p-value (t-test) (Column F), and 5) Storey q-value (Column G). A positive difference between mean normalized expressions indicate CM-specific over-expression, whereas a negative difference between mean normalized expressions indicate EPDC-specific over-expression.

**Table S11. Description and supporting literature of 91 signature genes in iPSC**

In this table, we describe the known functions of the 91 signature genes identified as differentially expressed between iPSC samples that differentiated to CMs and iPSC samples that differentiated into EDPCs (Table S14). For each gene, gene name (Column A), ensemble gene id (Column B), Chromosome (column C), gene start

(Column D) and end (Column E), the difference between mean normalized expressions (column F), the p-value (t-test) (Column G), and Storey q-value (Column H) are given. Additionally, for each gene functional descriptions (Column J) and PMIDs for supporting literature (Column I) are provided. This table is ordered on the difference between mean normalized expressions.

**Table S12: Regression estimates showing the associations between signature genes and %CM populations**

For each of the 91 signature genes (Columns A, B), linear regression estimate (Column C) and standard error (Column D) are shown. P-values (Column E) were calculated in R as 2*pnorm(estimate / standard error).

**Table S13. Independent contributions of 91 signature genes to cell fate determination**

For each of the 91 signature genes (Columns A, B), the $R^2$ is given (Column C).

**Table S14. Cumulative contribution of 91 signature genes to cell fate determination**

For each of the 35 signature genes (Columns A, B) that L1 norm identified as having significant contribution to cell fate determination, LASSO regression coefficient (Column C), median TPM (Column D), median contribution (Column E), and absolute value of the median contribution to the model (Column F) are shown.

**Table S15. Associations between genetic variation and differentiation outcome**

For each of 1,205 variants associated with the 91 signature genes in GTEx or iPSCs, shown are their chromosome (Column A), coordinates (Column B), reference and alternative allele (Columns C, D), linear regression estimate (Column E), standard error (Column F), p-value (Column G), and Storey q-value (Column H). P-values (Column I) were calculated in R as 2*pnorm (estimate / standard error).

**Table S16: GSEA showing functional enrichment between iPSCs that differentiated to CMs and iPSCs that differentiated to EPDCs**

For each of 9,808 MSigDB gene sets (Column A), GSEA enrichment (Column B), and p-value (Column C) calculated using the R gage package are shown. Storey q-value was used to adjust for multiple testing hypothesis. The analysis (Column E) shows whether the test was performed on all iPSCs (“All iPSC-CVPC samples”) or just on female samples (“Female samples”). Positive GSEA enrichment indicate enrichment for CM-specific gene sets, whereas negative GSEA enrichment indicate enrichment for EPDC-specific gene sets.

**Table S17. Table describing results of linear regression analysis to predict factors influencing differentiation potential of iPSC towards CM or EPDC fates**

Factors (Column A) input into the linear regression model. Columns B-E describe the results of the model, including estimate (Column B), standard error (Column C), z-value (Column D), and p-value (Column E).

**Table S18. Allelic imbalance fraction of genes on the X chromosome not in pseudoautosomal regions in females from iPSC samples and from iPSC-CVPC samples**

Gene_id indicates the ensemble gene id (Column A) for X chromosomes genes not in pseaudoautomsomal regions. Columns B-HR show the rna_assay_uuid (Table S5) of each of the female iPSC and iPSC-CVPC samples for which the allelic imbalance fraction was calculated for each gene.

**Table S19. Table describing differentiation outcomes and molecular data ID references from the Yoruba set**

Data from 39 Yoruba iPSC samples [14] (Column A) and their sex (Column B) are given. Outcome (Column C) indicates if the iPSC-CM differentiation was completed or terminated before completion of differenitation. %cTnT values (Column E), GEO iPSC RNA-seq sample IDs (Column F), and GEO iPSC-CM sample IDs (Column G) (GEO; GSE89895) are given.

**Table S20. Table describing observed beat scores and structure scores for iPSC-CVPC differentiations**

UDID (Column A) for each differentiation measured is given. Beat.Score (Column B) indicates the estimated beat score for the differentiation and Structure.Score indicates the observed structure score for the sample (Column C).

### REFERENCES


1. Liberzon, A. *et al.* Molecular signatures database (MSigDB) 3.0. *Bioinformatics* **27**, 1739-1740 (2011).
2. Subramanian, A. *et al.* Gene set enrichment analysis: a knowledge-based approach for interpreting genome-wide expression profiles. *Proceedings of the National Academy of Sciences of the United States of America* **102**, 15545-15550 (2005).
3. Lu, P., Takai, K., Weaver, V.M. & Werb, Z. Extracellular matrix degradation and remodeling in development and disease. *Cold Spring Harb Perspect Biol* **3** (2011).
4. Burridge, P.W. *et al.* Chemically defined generation of human cardiomyocytes. *Nature methods* **11**, 855-860 (2014).
5. Lian, X. *et al.* Robust cardiomyocyte differentiation from human pluripotent stem cells via temporal modulation of canonical Wnt signaling. *Proceedings of the National Academy of Sciences of the United States of America* **109**, E1848-1857 (2012).
6. Lian, X. *et al.* Directed cardiomyocyte differentiation from human pluripotent stem cells by modulating Wnt/beta-catenin signaling under fully defined conditions. *Nature protocols* **8**, 162-175 (2013).
7. Carrel, L. & Willard, H.F. X-inactivation profile reveals extensive variability in X-linked gene expression in females. *Nature* **434**, 400-404 (2005).
8. Tukiainen, T. *et al.* Landscape of X chromosome inactivation across human tissues. *Nature* **550**, 244-248 (2017).
9. Mekhoubad, S. *et al.* Erosion of dosage compensation impacts human iPSC disease modeling. *Cell stem cell* **10**, 595-609 (2012).
10. Patel, S. *et al.* Human Embryonic Stem Cells Do Not Change Their X Inactivation Status during Differentiation. *Cell Rep* **18**, 54-67 (2017).
11. Kadari, A. *et al.* Robust Generation of Cardiomyocytes from Human iPS Cells Requires Precise Modulation of BMP and WNT Signaling. *Stem cell reviews* **11**, 560-569 (2015).
12. Tohyama, S. *et al.* Distinct metabolic flow enables large-scale purification of mouse and human pluripotent stem cell-derived cardiomyocytes. *Cell stem cell* **12**, 127-137 (2013).
13. Panopoulos, A.D. *et al.* iPSCORE: A Resource of 222 iPSC Lines Enabling Functional Characterization of Genetic Variation across a Variety of Cell Types. *Stem Cell Reports* **8**, 1086-1100 (2017).
14. Banovich, N.E. *et al.* Impact of regulatory variation across human iPSCs and differentiated cells. *Genome research* **28**, 122-131 (2018).
